# Transcriptomic correlates of neuron electrophysiological diversity

**DOI:** 10.1101/134791

**Authors:** Shreejoy J. Tripathy, Lilah Toker, Brenna Li, Cindy-Lee Crichlow, Dmitry Tebaykin, B. Ogan Mancarci, Paul Pavlidis

## Abstract

How neuronal diversity emerges from complex patterns of gene expression remains poorly understood. Here we present an approach to understand electrophysiological diversity through gene expression by integrating transcriptomics with intracellular electrophysiology. Using a brain-wide dataset of 34 neuron types, we identified 420 genes whose expression levels significantly correlated with variability in one or more of 11 electrophysiological parameters. The majority of these correlations were consistent in an independent sample of 12 visual cortex cell types. Many associations reported here have the potential to provide new insights into how neurons generate functional diversity, and correlations of ion channel genes like *Gabrd* and *Scn1a* (Nav1.1) with resting potential and spiking frequency are consistent with known causal mechanisms. These results suggest that despite the complexity linking gene expression to electrophysiology, there are likely some general principles that govern how individual genes establish phenotypic diversity across very different cell types.

## Introduction

A major goal of neuroscience has been to understand the mechanistic origins of neuronal electrophysiological phenotypes. Such electrical features help define the computational functions of each neuron ^1,2^, and further, specific electrophysiological deficits contribute to brain disorders such as epilepsy, ataxia, and autism ^3–5^.

The molecular basis of neuron electrophysiology is complex. There are over 200 mammalian ion channel and transporter genes whose products influence a neuron’s electrophysiological phenotype ^6–9^. Numerous additional genes regulate channel functional expression through initiating gene transcription and alternative splicing, post-translational modifications, and trafficking channels to and from the membrane surface ^10–12^. Even morphological features contribute to cellular electrophysiology ^13^. Recent genetic studies in human epileptic and neuropsychiatric patients provide convergent evidence, as mutations in many genes reflecting multiple functional pathways are associated with these disorders ^4,14–16^. In light of this complexity, the gold standard employed by neurophysiologists is to use gene knockouts or pharmacology to assay how electrophysiological function changes following protein disruption ^7,8^. However, these single-gene focused methods are relatively low-throughput and many potentially relevant genes have yet to be studied for their electrophysiological function.

Cell type-specific transcriptomics, enabling genome-wide assay of quantitative mRNA expression levels, provides a lucrative avenue for discovering novel genes that might contribute to specific aspects of cellular physiology ^17,18^. Correlation-based approaches have been proposed that pair single-cell expression profiling with patch-clamp electrophysiology ^19–21^. These approaches leverage the biological variability observed across a collection of cells to identify gene expression patterns correlated with cellular phenotypic differences. However, generalizing from these studies has proven challenging however, since they typically have been focused on a limited number of cell types. Similarly, and perhaps more critically, there are typically hundreds to thousands of genes correlated with electrophysiological variability^22^. Thus it has been difficult from this data to pin down how individual genes might shape specific cellular phenotypes. Though making use of larger and more diverse collections of cell types could provide a potential solution, collecting such reference data is immensely resource- and labor-intensive.

Here, we present an approach for correlating cell type-specific transcriptomics with neuronal electrophysiological features. We employ a novel reference dataset on brain-wide neuronal gene expression and electrophysiological diversity, reflecting the neuronal characterization efforts of hundreds of investigators as well as our recent work to compile and standardize these data ^23–25^. We identified hundreds of genes whose expression levels significantly correlate with specific electrophysiological features (e.g., resting potential or maximum spiking frequency). Illustrating the generalizability of these results, most correlations were also consistent with an independent neocortex-specific dataset from the Allen Institute. Many of these genes have been further found to directly regulate neuronal electrophysiology, suggesting that some of the correlations reported here likely reflect novel causal relationships. Our findings present a major step for understanding how a multitude of genes contribute to cell type-specific phenotypic diversity.

## Results

Our overall approach was to first compile a reference dataset of brain cell type-specific transcriptomes paired with cell type-specific electrophysiological (ephys) profiles. We then assessed the ability of gene expression to statistically explain variance in specific ephys properties. We next validated whether these gene-ephys correlations were consistent with an independent dataset on visual cortex neurons collected by the Allen Institute for Brain Science (AIBS). Lastly we made use of literature review to establish whether any of these gene-ephys correlations had been previously shown to be causal.

### Discovery and validation datasets

To construct our primary dataset for gene-ephys correlation analysis, we combined two databases developed and curated by our group. The first, NeuroExpresso, a database containing microarray-based transcriptomes collected from samples of purified mouse brain cell types under normal conditions 23. The second, NeuroElectro, a database of neuronal electrophysiological profiles manually curated from the published literature on rodent intracellular electrode recordings from normal, non-treated cells,^24,25^. Given the methodological heterogeneity of the primary data comprising these databases, we applied a number of quality control filtering and cross-laboratory standardization and normalization methods (see Methods). We obtained neuron type-specific gene expression and ephys data by merging these databases on cell type identity, making use of our detailed annotations of each sample’s specific cell type (Figure 1A). The final “discovery” reference dataset is composed of 34 neuron types sampled throughout the brain and reflects cell types with diverse circuit roles, neurotransmitters, and developmental stages (summarized in Table 1 and Supplemental Table 2).

**Figure 1.**
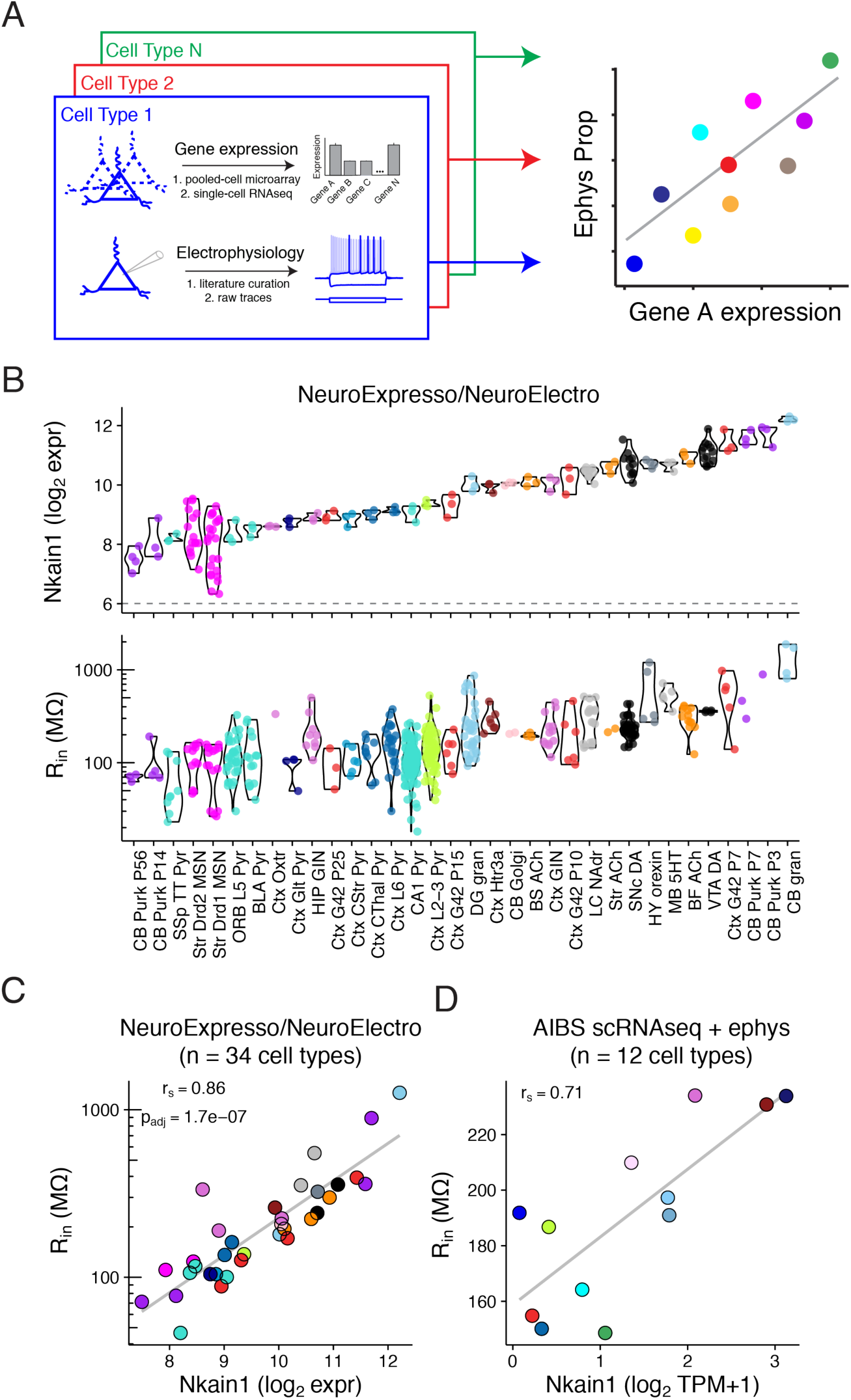
Correlating cell type-specific gene expression with electrophysiological diversity. A) Illustration of transcriptomic and ephys data compilation by cell type (left) and correlation analysis of single gene expression by ephys parameter diversity (right). B) Top row: Gene expression levels of *Nkain1* across 34 neuron types sampled from the combined NeuroExpresso/NeuroElectro dataset. Each dot reflects a unique transcriptomic sample collected from purified cells and y-axis is in units of log2 expression (i.e., each increment reflects a 2-fold change in expression level). Dashed line at 6 indicates approximate level of background expression. Bottom row: Input resistance values for the same cell types in top row. Individual dots reflect population mean electrophysiological values manually curated from individual articles represented in the NeuroElectro database, following experimental condition normalization. C) Same data as in B, but data has been summarized by the mean (expression, x-axis) or median (ephys, y-axis) value within each cell type. r_s_ indicates Spearman rank correlation and p_adj_ indicates Benjamini Hochberg false discovery rate. Note that cell types with high *R*_*in*_, such as cerebellar granule cells and midbrain dopaminergic cells, express high levels of *Nkain1* whereas cell types with low *R*_*in*_, including neocortical and hippocampal pyramidal cells, express low levels of *Nkain1*. D) Corresponding summary data from the Allen Institute for Brain Science (AIBS) Cell Types dataset. Dots reflect averaged values from 12 individual mouse cre-lines and are detailed in *Table 2*. Expression values are based on single-cell RNAseq (scRNAseq), quantified as Transcripts Per Million (TPM). Ephys values are based on single-cell characterization *in vitro*.

**Table 1:**
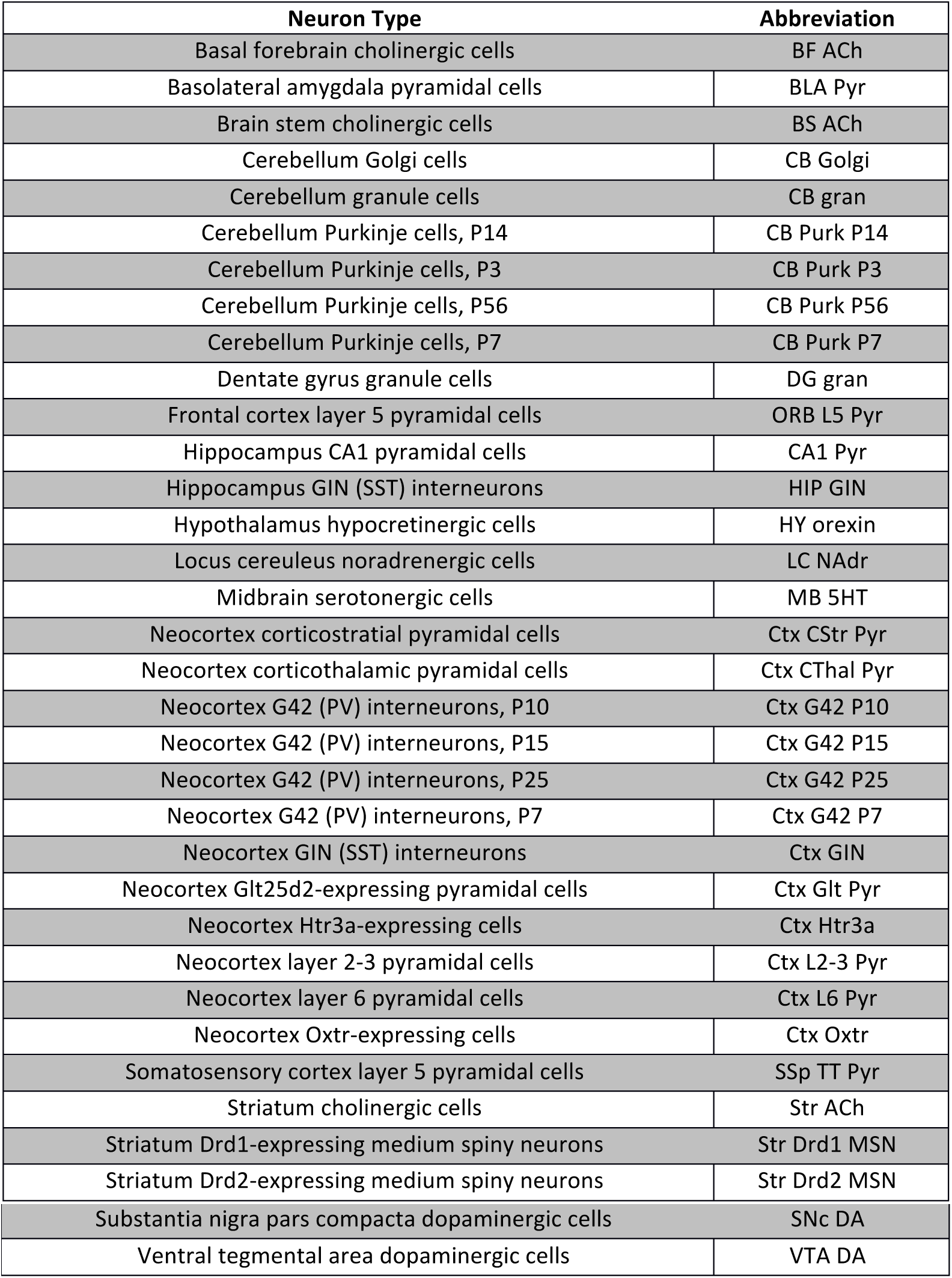
Descriptions for neuron types composing the NeuroExpresso/NeuroElectro discovery dataset. References for individual transcriptomic and electrophysiological samples are available in *Supplemental Table 2*.

| Neuron Type | Abbreviation |
| --- | --- |
| Basal forebrain cholinergic cells | BF ACh |
| Basolateral amygdala pyramidal cells | BLA Pyr |
| Brain stem cholinergic cells | BS ACh |
| Cerebellum Golgi cells | CB Golgi |
| Cerebellum granule cells | CB gran |
| Cerebellum Purkinje cells, P14 | CB Purk P14 |
| Cerebellum Purkinje cells, P3 | CB Purk P3 |
| Cerebellum Purkinje cells, P56 | CB Purk P56 |
| Cerebellum Purkinje cells, P7 | CB Purk P7 |
| Dentate gyrus granule cells | DG gran |
| Frontal cortex layer 5 pyramidal cells | ORB L5 Pyr |
| Hippocampus CA1 pyramidal cells | CA1 Pyr |
| Hippocampus GIN (SST) interneurons | HIP GIN |
| Hypothalamus hypocretinergic cells | HY orexin |
| Locus cereuleus noradrenergic cells | LC NAdr |
| Midbrain serotonergic cells | MB 5HT |
| Neocortex corticostratial pyramidal cells | Ctx CStr Pyr |
| Neocortex corticothalamic pyramidal cells | Ctx CThal Pyr |
| Neocortex G42 (PV) interneurons, P10 | Ctx G42 P10 |
| Neocortex G42 (PV) interneurons, P15 | Ctx G42 P15 |
| Neocortex G42 (PV) interneurons, P25 | Ctx G42 P25 |
| Neocortex G42 (PV) interneurons, P7 | Ctx G42 P7 |
| Neocortex GIN (SST) interneurons | Ctx GIN |
| Neocortex Glt25d2-expressing pyramidal cells | Ctx Glt Pyr |
| Neocortex Htr3a-expressing cells | Ctx Htr3a |
| Neocortex layer 2-3 pyramidal cells | Ctx L2-3 Pyr |
| Neocortex layer 6 pyramidal cells | Ctx L6 Pyr |
| Neocortex Oxtr-expressing cells | Ctx Oxtr |
| Somatosensory cortex layer 5 pyramidal cells | SSp TT Pyr |
| Striatum cholinergic cells | Str ACh |
| Striatum Drd1-expressing medium spiny neurons | Str Drd1 MSN |
| Striatum Drd2-expressing medium spiny neurons | Str Drd2 MSN |
| Substantia nigra pars compacta dopaminergic cells | SNc DA |
| Ventral tegmental area dopaminergic cells | VTA DA |

For validation we utilized an independent dataset characterizing neurons from adult mouse primary visual cortex collected by the Allen Institute for Brain Science. Here, genetically labeled cells were characterized either for their transcriptomic profiles, using single-cell RNA sequencing (scRNAseq) ^26^, or their electrophysiological properties, using patch-clamp electrophysiology *in vitro* with standardized protocols (http://celltypes.brain-map.org/). Importantly, for both expression and ephys characterization, the same mouse lines for genetically labeling specific populations of cells were used, making it straightforward to combine samples post-hoc, yielding a final “validation” dataset composed of 12 unique cell types (Table 2). Averaging data across labeled single cells within a mouse line also helps mitigate the influence of cell-to-cell variability and technical “dropouts” in the scRNAseq data ^18^. Given the smaller number of cell types present in the AIBS dataset we chose to use these data primarily for validation and generalization of findings made using discovery dataset. Note that for both the discovery and validation datasets, electrophysiological and gene expression values are from separate cells.

**Table 2:** Descriptions for neuron types composing the Allen Institutes for Brain Sciences Cell Types validation dataset. Mouse line indicates cre-driver lines used to label specific populations of cells in the adult mouse visual cortex. N cells indicates number of cells assayed per cre-line via single-cell RNAseq or patch-clamp electrophysiology. Color indicates cell type color used within this manuscript.

| Mouse line (cre-driver) | N cells (scRNAseq) | N cells (ephys) | Color |
| --- | --- | --- | --- |
| **Ctgf** | 23 | 12 | midnightblue |
| **Cux2** | 122 | 55 | olivedrab1 |
| **Gad2** | 77 | 11 | thistle1 |
| **Htr3a** | 123 | 81 | firebrick4 |
| **Nr5a1** | 48 | 62 | blue2 |
| **Ntsr1** | 90 | 37 | deepskyblue |
| **Pvalb** | 89 | 141 | firebrick2 |
| **Rbp4** | 173 | 61 | mediumseagreen |
| **Rorb** | 54 | 106 | skyblue3 |
| **Scnn1a.Tg2** | 19 | 28 | cyan |
| **Scnn1a.Tg3** | 99 | 52 | lightskyblue |
| **Sst** | 107 | 107 | orchid |

### Analysis approach

Our primary analysis focus was to understand how cell type-specific expression of individual genes might statistically explain the variance in electrophysiological parameters observed across cell types (Figure 1A, right). For example, how does *Scn1a* (Na_v_1.1) expression correlate with neuronal maximum firing rates? which genes are most correlated with cellular resting membrane potentials? We primarily chose to employ a single-gene focused approach because of sample size considerations, reasoning that we did not have enough unique cell types in either the discovery or validation datasets to rigorously pursue a combinatorial gene approach. However, this single-gene focus might limit our ability to identify highly combinatorial and/or redundant or degenerate relationships between gene expression profiles and ephys ^27,28^.

### Correlation of neuronal transcriptomics with electrophysiological properties

For each of the 34 neuron types in the NeuroExpresso/NeuroElectro discovery dataset, we obtained a gene expression profile for 11,509 genes and 5-11 intrinsic electrophysiological properties (mean = 9 +/- 2 ephys properties per cell type; described in Supplemental Table 1). We first asked whether there are individual genes whose quantitative mRNA expression levels correlate with systematic ephys diversity in the both the discovery and AIBS validation datasets. Using the discovery dataset, after first filtering for genes with sufficiently high and variable expression across cell types (see Methods), we found a total of 653 genes (of 2694 tested) correlated with at least 1 of the 11 ephys properties at p_adj_ < 0.05 (p_adj_ indicates Benjamini-Hochberg false discovery rate adjusted p-value). 1095 genes were identified at p_adj_ < 0.1 and 217 genes were identified at p_adj_ < 0.01.

As an illustrative example of one gene-ephys correlation, we found that expression levels of the gene *Nkain1* correlated with input resistance (R_in_) values across cell types in the discovery dataset (Figure 1B, C; Spearman correlation, r_s_ = 0.86; p_adj_ = 1.7*10^-7^). We also saw this trend recapitulated when only considering within-cell type changes observed across cortical basket cell and Purkinje cell development, with *Nkain1* expression and R_in_ decreasing dramatically as these cells mature (Supplemental Figure 1). In the AIBS validation dataset, after summarizing the single-cell data to the level of cell types, we further found a consistent *Nkain1-* R_in_ correlation amongst adult visual cortex cell types (Figure 1D; r_s_ = 0.71). Little is known about *Nkain1* protein function, except that it interacts with the Na^+^/K^+^ pump β-subunit and likely modulates the pump’s function and membrane localization ^29^. Intriguingly, the Na+/K+ pump has a known role in establishing cellular volumes and input resistance ^30^.

**Supplemental Figure 1.**
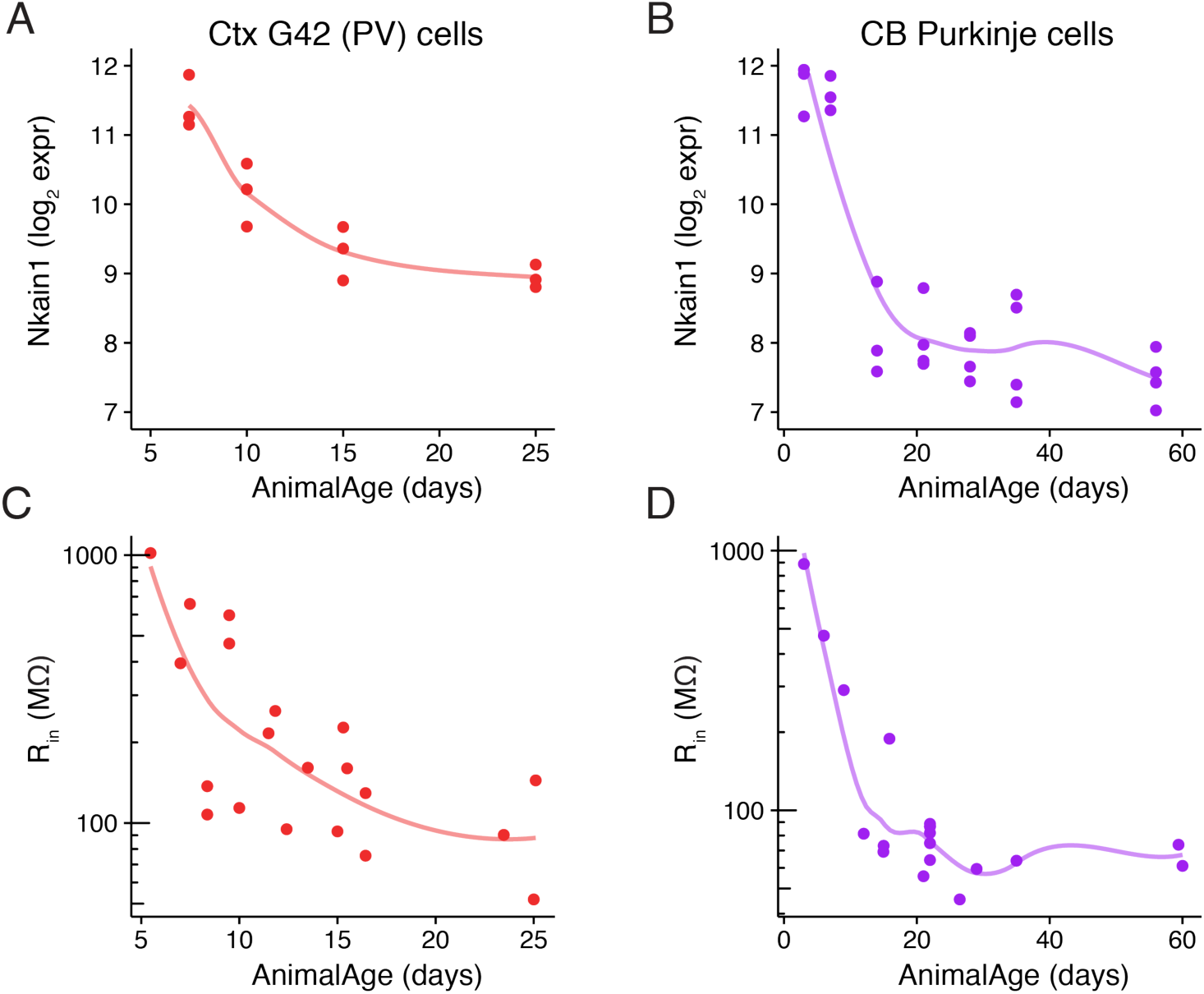
Example of cell type-specific transcriptomic and electrophysiological changes across development. A) Gene expression levels of *Nkain1* across development of cortical G42 parvalbumin-expressing interneurons. Dots reflect unique transcriptomic samples. B) Same as A, but for cerebellar Purkinje cells. C) Values of input resistance sampled from cortical G42 parvalbumin-expressing interneurons at various points in development. Individual dots reflect population means from individual articles represented in the NeuroElectro database and lines are based on loess smoothing. D) Same as C, but for cerebellar Purkinje cells.

We provide a summary of the total number of genes identified as significantly correlated with each of the 11 ephys properties in Figure 2A and the full list of gene-ephys correlations in Supplemental Table 3. We initially noticed that different ephys properties were significantly correlated with varying numbers of genes. For example, at the somewhat conservative threshold of p_adj_ < 0.05, we found no genes correlated with action potential threshold voltage (AP_thr_), despite there being many genes previously implicated with this feature ^5,31^. In contrast, there were over 200 genes significantly correlated with either V_rest_ or AHP_amp_. However, we consider it unlikely that all of these genes reflect a direct causal relationship, as gene-gene correlations driven by gene co-regulation create ambiguity.

**Figure 2:**
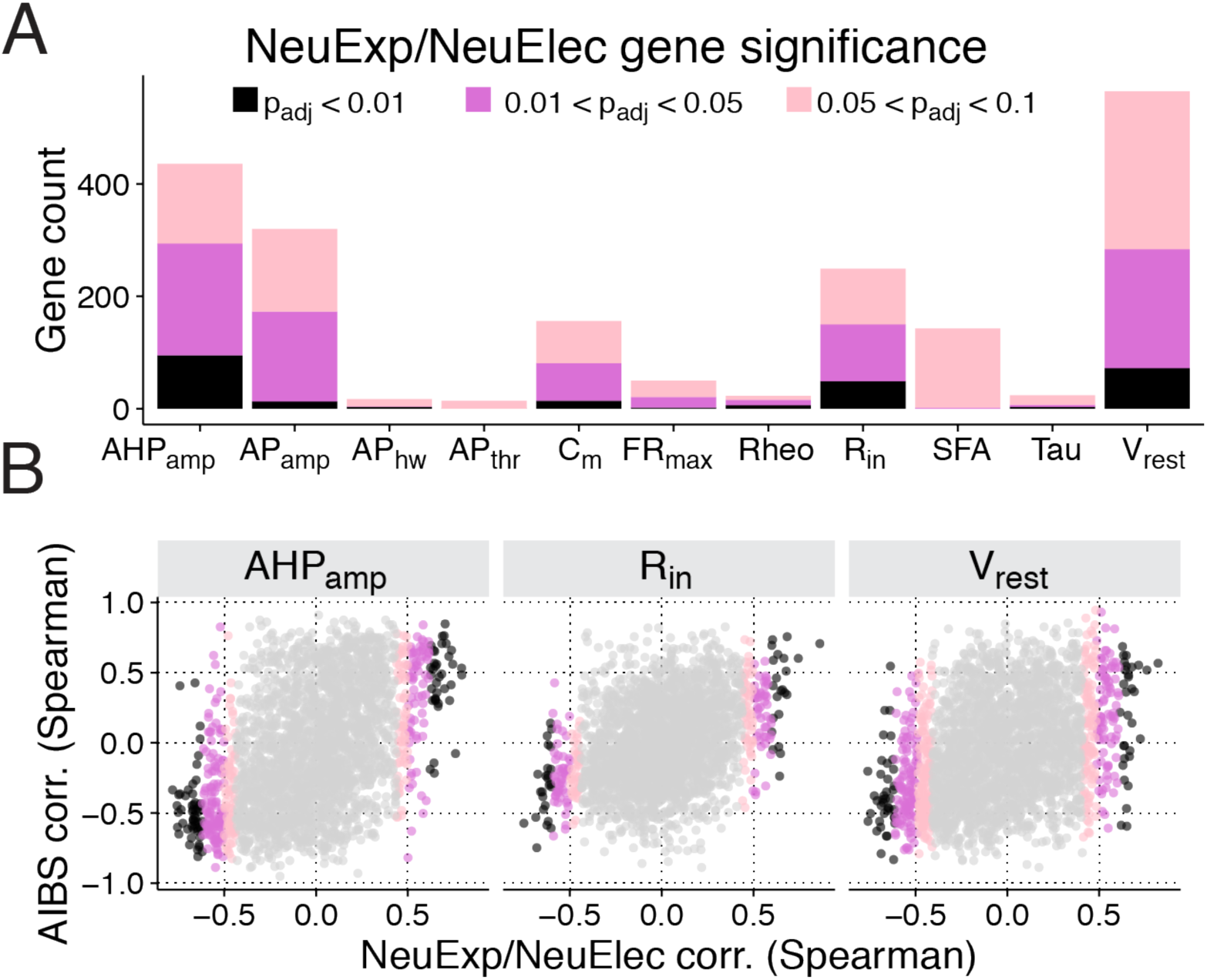
Identification and validation of transcriptomic - electrophysiological correlations. A) Count of genes significantly correlated with various electrophysiological properties, broken down by statistical significance of Benjamini-Hochberg FDR-adjusted correlation p-values (p_adj_). Names and descriptions of ephys properties are provided in *Supplemental Table 1*. B) Comparison of correlations calculated using NeuroExpresso/ NeuroElectro discovery dataset (NeuExp/NeuElec, x-axis) versus correlations calculated using Allen Institute validation dataset (AIBS, y-axis). Dots reflect correlation values of individual genes. Subpanels indicate correlations computed across various electrophysiological properties and p-values are provided in *Table 3*.

We note that in the discovery dataset, not all ephys properties were available for each cell type, with 19-34 cell types quantified per ephys property. Furthermore, since correlation p-values are in part related to sample size, we found a positive relationship between the total number of genes associated with each ephys property and the number of cell types where the ephys property was quantified (R^2^ = 0.30; Supplemental Figure 2). Next, given that ephys properties tend to be correlated with one another ^21,25^, we asked if pairs of correlated ephys properties also tend to share associated genes. For example, cellular measurements of membrane capacitance (C_m_) and R_in_ are highly anti-correlated (r_s_ = -0.69 in the discovery dataset); furthermore, of the 80 genes significantly associated with C_m_, 36 were also associated with R_in_. Though some pairs of ephys properties share common biophysical mechanisms and could be thus regulated via common genes (e.g., C_m_ and R_in_ are both dependent in part on cell size), correlations between ephys properties likely limit the specificity of the relationships reported here.

We next used the AIBS dataset to validate the significant correlations observed in the discovery dataset. We predicted that gene-ephys correlations discovered in our brain-wide dataset should generalize to the transcriptomic and electrophysiological diversity among adult visual cortex cell types. Because of the limited number of cell types available in the AIBS dataset, we compared results between the discovery and validation datasets as: 1) overall consistency, defined by the global rank correlation between results from the two datasets (Figure 2B); and 2) consistency for the subset of gene-ephys relationships meeting our threshold for significance in the discovery dataset (p_adj_ < 0.05). Overall, we found positive, but modest, agreement between the two datasets, with most ephys properties showing a positive correlation (Table 3). However, AP_thr_, Rheo, and Tau are notable exceptions and might reflect challenges in normalizing these ephys features from the cross-study NeuroElectro database ^25^. Focusing specifically on significant gene-ephys correlations identified in the discovery dataset, we found that the majority of these, 61.2%, reflecting 420 individual genes, were consistent in the validation dataset, with consistency defined as a matching correlation direction and with an absolute value of r_s_ > 0.3 (Table 3).

**Table 3:** Consistency of gene-electrophysiological property correlations between NeuroExpresso/NeuroElectro discovery and AIBS validation datasets. Overall AIBS consistency indicates overall rank correlation between gene-electrophysiological correlations calculated in both the discovery and validation datasets. P-values based on 1000 random reshuffles of cell type labels in the AIBS validation dataset. Discovered genes, *p*_*adj*_ < 0.05 reflects count of genes significantly correlated with each ephys property with in discovery dataset (only includes genes that are also present in AIBS scRNAseq dataset). AIBS consistency, |r_s_|> 0.3 reflects count and percentage of discovered genes that further show a consistent relationship in the AIBS validation dataset. P-value also based on 1000 shuffled samples of cell type labels in the validation dataset.

| Ephys Property | Overall AIBS consistency | | Discovered genes;<br>$p_{\text{adj}} < 0.05$ | AIBS consistency;<br>$\vert r_s\vert > 0.3$ | | |
| --- | --- | --- | --- | --- | --- | --- |
|  | *Spearman corr.* | *p-value* | *count* | *count* | *%* | *p-value* |
| **AHP<sub>amp</sub>** | 0.45 | 0.009 | 285 | 204 | 72 | 0.005 |
| **AP<sub>amp</sub>** | 0.404 | <0.001 | 169 | 119 | 70 | 0.006 |
| **AP<sub>hw</sub>** | 0.04 | 0.323 | 4 | 3 | 75 | 0.056 |
| **AP<sub>thr</sub>** | -0.146 | 0.877 | 0 | --- | --- | --- |
| **C<sub>m</sub>** | 0.384 | 0.037 | 80 | 55 | 69 | 0.015 |
| **FR<sub>max</sub>** | 0.209 | 0.074 | 21 | 7 | 33 | 0.159 |
| **R<sub>heo</sub>** | -0.049 | 0.649 | 15 | 5 | 33 | 0.162 |
| **R<sub>in</sub>** | 0.346 | 0.004 | 144 | 68 | 47 | 0.029 |
| **SFA** | 0.298 | 0.01 | 2 | 1 | 50 | 0.277 |
| **Tau** | -0.106 | 0.713 | 6 | 5 | 83 | 0.007 |
| $V_{rest}$ | 0.332 | 0.029 | 279 | 148 | 53 | 0.025 |

The degree of consistency between the NeuroExpresso/NeuroElectro and AIBS datasets is encouraging given their dissimilarity in design and content. For example, the AIBS cell types dataset is sampled from a single brain region (visual cortex) at one developmental stage (adult). Moreover, there are considerable technical differences between the datasets, such as transcriptome quantification via single-cell RNAseq vs pooled-cell microarrays or between standardized versus heterogeneous ephys data collection.

In the remainder of the manuscript, we focus on further characterizing the significant gene-ephys correlations from the discovery dataset that have evidence for further validating in the AIBS dataset.

**Supplemental Figure 2:**
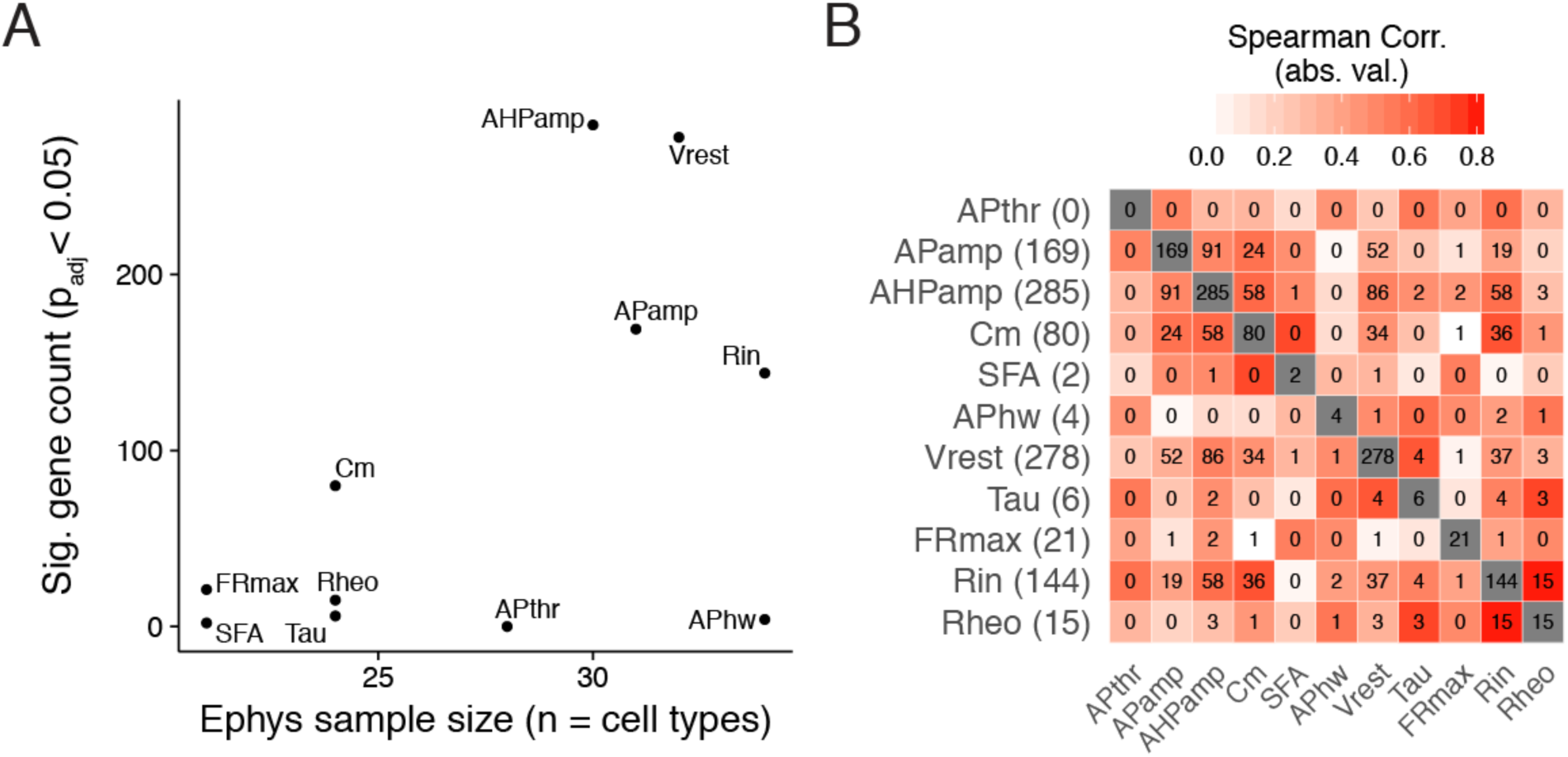
Factors affecting numbers of genes identified as significantly correlated with different electrophysiological properties. A) Scatterplot illustrating the relationship between the numbers of genes identified as significantly correlated with each ephys property (p_adj_ < 0.05) versus the number of cell types with ephys data in the NeuroExpresso/NeuroElectro dataset. B) Pairwise correlations between electrophysiological properties, based on cell types in combined NeuroExpresso/NeuroElectro sample. Heatmap colors indicate the absolute value of measured Spearman correlations between ephys property pairs. Inset values indicate the number of significant genes shared between each pair of ephys properties (p_adj_ < 0.05). Numbers in parentheses on y-axis and values along diagonal indicate number of significant genes identified for each ephys property (i.e., as in y-axis in A).

### Causal relationships between discovered gene-electrophysiological correlations

A key question is whether any of the gene-ephys correlations we observed are due to direct causal relationships supported by specific evidence. To this end, we made use of the existing literature on gene-ephys relations. We focused on ion channel genes, reasoning that these would be most likely to have been directly tested for electrophysiological function. We manually searched the literature for such experiments, since at present this data is not reflected within a comprehensive database (the current NeuroElectro database reflects experiments done under standard or control conditions, not genetic or pharmacological manipulations).

We present a brief summary of our gene-centered literature search alongside highlights from our correlation-based analysis below, with the complete results provided in Supplemental Table 4. Of 31 significant and validated ion channel-ephys correlations, we found 17 had been directly tested through genetic manipulations or channel-specific pharmacology (reflecting 12 unique ion channel genes). To compare our correlations to individual results from direct experiments, we first mapped our correlations to predicted causal effects; for example, knocking out a gene whose expression is positively correlated with maximum firing rate should tend to lower firing rates, all else being equal. We found that of 17 total tested ion channel-ephys correlations; 11 were consistent with literature evidence, 2 showed mixed evidence, 1 showed no effect on the ephys property, and 3 were inconsistent. Here, we defined inconsistent evidence as those where a predicted increase (or decrease) in an ephys property was reflected by a change in the opposite direction in the literature; mixed evidence were those where some manipulations were consistent but others were inconsistent (e.g., pharmacology versus gene knockout). Below, we provide specific illustrative examples from this literature search.

*Scn1a*, encoding the sodium channel Nav1.1, was positively correlated with maximum firing rate (Figure 3B; NeuExp/NeuElec r_s_ = 0.86, AIBS r_s_ = 0.36), with the highest *Scn1a* expression observed in adult cortical PV interneurons and Purkinje cells. In a mouse model of Dravet syndrome with a hemizygous gene deletion (i.e., *Scn1a* +/-), it was observed that fast-spiking PV interneurons cells could no longer fire at their characteristically high frequencies (Figure 3C), with a smaller but significant effect also observed in Sst-expressing Martinotti cells ^5^. However, the same change was not seen in layer 5 pyramidal cells, which express ˜3–4 fold less *Scn1a* relative to PV cells (in NeuroExpresso and AIBS), potentially suggesting that total expression levels might mediate the effect of hemizygous *Scn1a* deletion. Intriguingly, in a haploinsufficiency model of Dravet syndrome, directly upregulating *Scn1a* expression using long non-coding RNAs rescued the firing phenotype in PV cells and lowered seizure number and duration ^32^.

**Figure 3:**
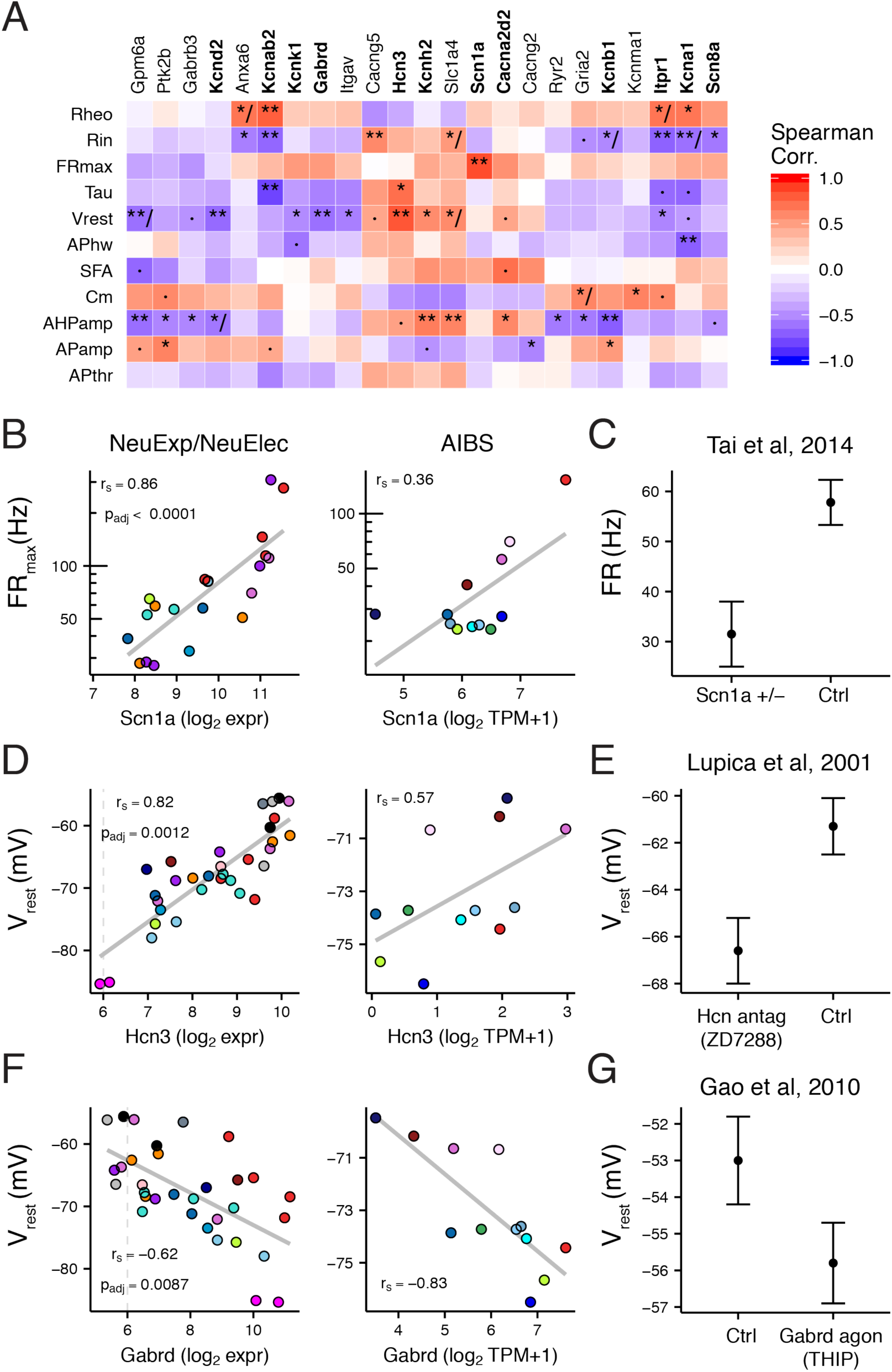
Ion channel specific gene-electrophysiological correlations and literature evidence for causal regulation. A) Heatmap showing NeuExp/NeuElec dataset gene-ephys correlations for ion channel genes. Genes filtered for those with at least one significant ephys correlation (p_adj_ < 0.05) and with validation supported in AIBS dataset. Gene names in bold indicate those we found to be previously studied for specific predicted ephys properties, based on our literature search. Symbols within heatmap: ·, p_adj_ < 0.1; *, p_adj_ < 0.05; **, p_adj_ < 0.01; /, indicates inconsistency between discovery and AIBS validation dataset. B) Correlation between cell type-specific *Scn1a* (Na_v_1.1) gene expression and maximum firing rate (FR_max_) from discovery dataset (NeuExp/NeuElec, left) and Allen Institute dataset (AIBS, right). Grey trend lines indicate linear fit. C) Replotted data from Tai et al 2014, showing evoked firing rates at 300 pA current injection for parvalbumin positive interneurons in control and Scn1a heterozygous mice (Scn1a +/-). Data plotted as mean +/- SEM. D) Same as B, but for *Hcn3* and resting membrane potential (V_rest_). E) Replotted data from Lupica et al 2001, where V_rest_ from CA1 OLM interneurons was measured before and after the application of ZD7288, a selective antagonist of HCN channels. F) Same as B, but for *Gabrd* and V_rest_. G) Replotted data from Gao et al, 2010, showing V_rest_ recorded from dorsal motor nucleus of vagus neurons after application of THIP, a selective agonist of *Gabrd*-mediated tonic inhibition.

We found 4 (of 5 total) ion channel genes correlated with V_rest_ that were consistent with literature evidence. *Hcn3*, encoding a slow HCN channel variant ^6^, was positively correlated with V_rest_ (Figure 3D; NeuExp/NeuElec r_s_ = 0.82, AIBS r_s_ = 0.57). Blocking HCN-current using ZD7288 across multiple cell types consistently made V_rest_ more hyperpolarized (Figure 3 E) ^33,34^. *Gabrd*, *Kcnk1*, and *Itpr1*, were each negatively correlated with V_rest_ and each gene reflects a different mechanistic route towards V_rest_ hyperpolarization (Figure 3F and Supplemental Figure 3). For example, *Gabrd* encodes the δ-subunit of the GABA_A_ receptor and mediates extrasynaptic tonic inhibition, effectively turning the GABA_A_ receptor into a chloride channel^35^. Thus, increased *Gabrd* expression, or pharmacologically increasing its activity (Figure 3F,G)^36^ would tend to hyperpolarize cells through the chloride reversal potential (median E_Cl_ = -72 mV, based on reported internal and external solutions). Similarly, *Kcnk1*, encoding the K_2P_1.1 2-pore potassium channel, hyperpolarizes V_rest_ through the potassium reversal potential (E_K_ ˜ -100 mV)^37^. *Itpr1* activity releases calcium from intracellular stores and hyperpolarizes V_rest_ through calcium-activated potassium channels ^38,39^. Taken together, each of these genes reflect distinct and potentially degnerate routes towards modulating cellular V_rest_.

We found evidence for two ion channel subunits, *Kcna1* and *Kcnab2*, regulating multiple distinct electrophysiological properties (Supplemental Figure 3). For example, *Kcna1*, encoding the delayed rectifier potassium channel Kv1.1, was negatively correlated with action potential half width (NeuExp/NeuElec r_s_ = -0.70, AIBS r_s_ = -0.52) and positively correlated with rheobase (NeuExp/NeuElec r_s_ = 0.69, AIBS r_s_ = 0.66). These correlations were corroborated by *Kcna1* genetic knockouts or pharmacological block in auditory brainstem neurons and are consistent with known mechanistic insight about Kv1.1 function ^40,41^.

While the previous examples are encouraging, not all of our findings were concordant with previous literature. For example, we saw that *Kcnb1*, encoding the Kv2.1 channel, was negatively correlated with spike afterhyperpolarization amplitude (AHP_amp_) (Supplemental Figure 4A;B; NeuExp/NeuElec r_s_ = -0.70, p_adj_ = 0.0033; AIBS r_s_ = -0.62). Based on this correlation, we would expect that decreasing K_v_2.1 functional expression should increase AHP_amp_ values. However, convergent genetic and pharmacological evidence suggests the opposite: decreasing K_v_2.1 activity or expression decreases AHP_amp_ values ^42,43^. Delving deeper, the *Kcnb1-AHP_amp_* correlation appears driven in part by gross differences between excitatory and non-excitatory cell types, with excitatory cells strongly expressing *Kcnb1* and also having small AHP_amp_ relative to non-excitatory cell types (Supplemental Figure 4C). Thus though there is likely some mechanistic explanation for why excitatory cells tend to express more *Kcnb1*, this does not appear to be directly related to AHP_amp_ per-se. This example suggests that caution is needed before interpreting each correlation reported here as a direct causal relationship.

To summarize, we found multiple examples of direct regulation of specific ephys properties by individual genes identified through our correlation-based methodology. In the remainder of the results, we highlight additional genes that may be of relevance in future studies.

**Supplemental Figure 3:**
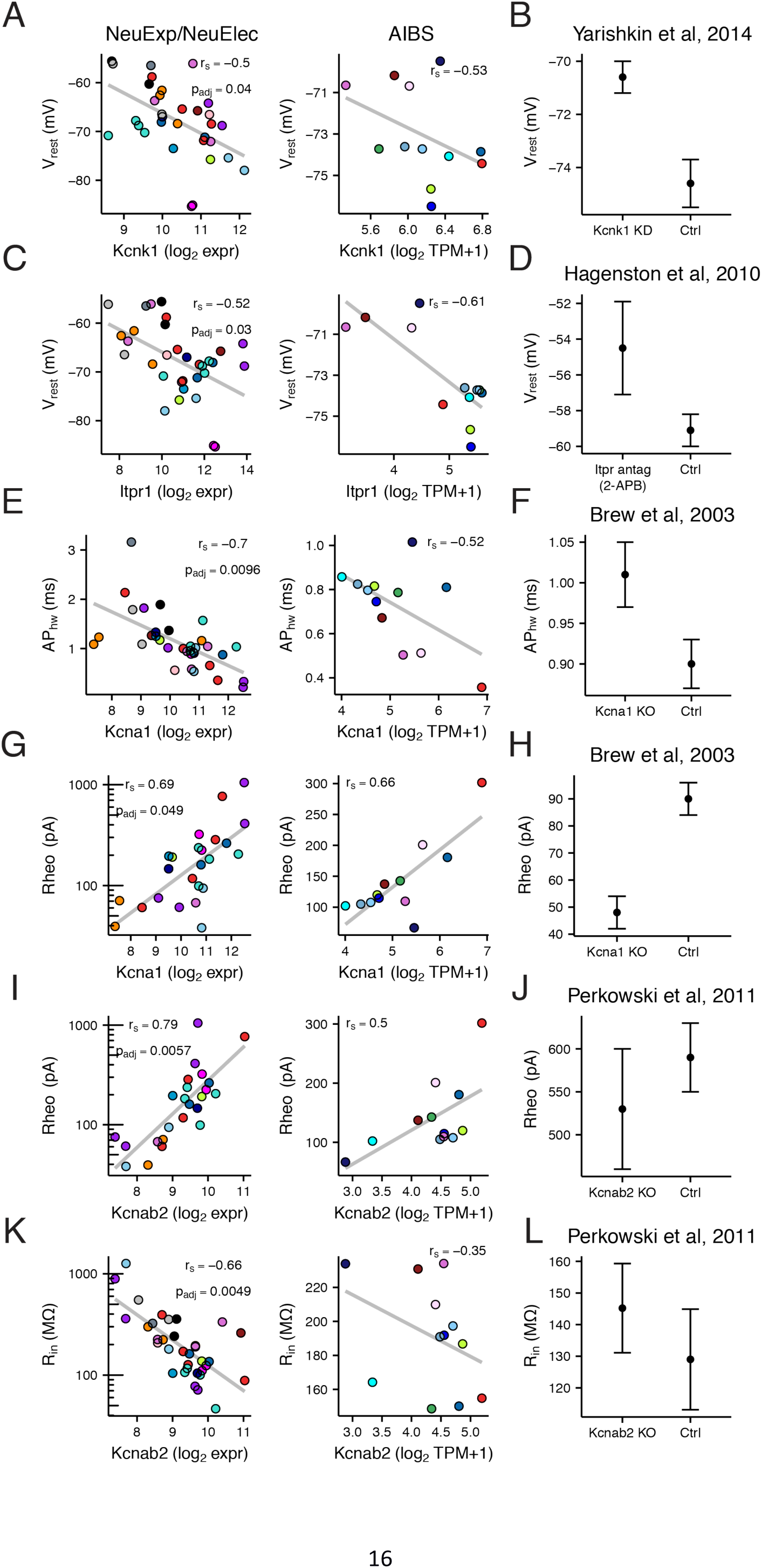
Further evidence for causal regulation of specific gene-ephys correlations. A) Correlation between cell type-specific *Kcnk1* (K_2P_1.1/TWIK1) gene expression and resting membrane potential (V_rest_) from discovery dataset (NeuExp/NeuElec, left) and Allen Institute dataset (AIBS, right). B) Replotted data from Yarishkin et al, 2014, showing effects of siRNA-induced knockdown of *Kcnk1* expression in dentate gyrus granule cells. C, E, I, G, K) Same as A but shown for specific ephys properties and genes. D) Replotted data from Hagenston et al, 2010, showing effects of antagonizing *Itpr1* function through the use of 2-APB. F, H) Replotted data from Brew et al, 2010, showing effects of knocking out *Kcna1* (K_v_1.1) on action potential half width (AP_hw_) and rheobase (Rheo) as measured in auditory brainstem neurons. J, L) Replotted data from Perkowski et al, 2011, showing effects of knocking out *Kcnab2* (K_v_beta2) on rheobase and input resistance (R_in_) as measured in lateral amygdala pyramidal neurons.

**Supplemental Figure 4:**
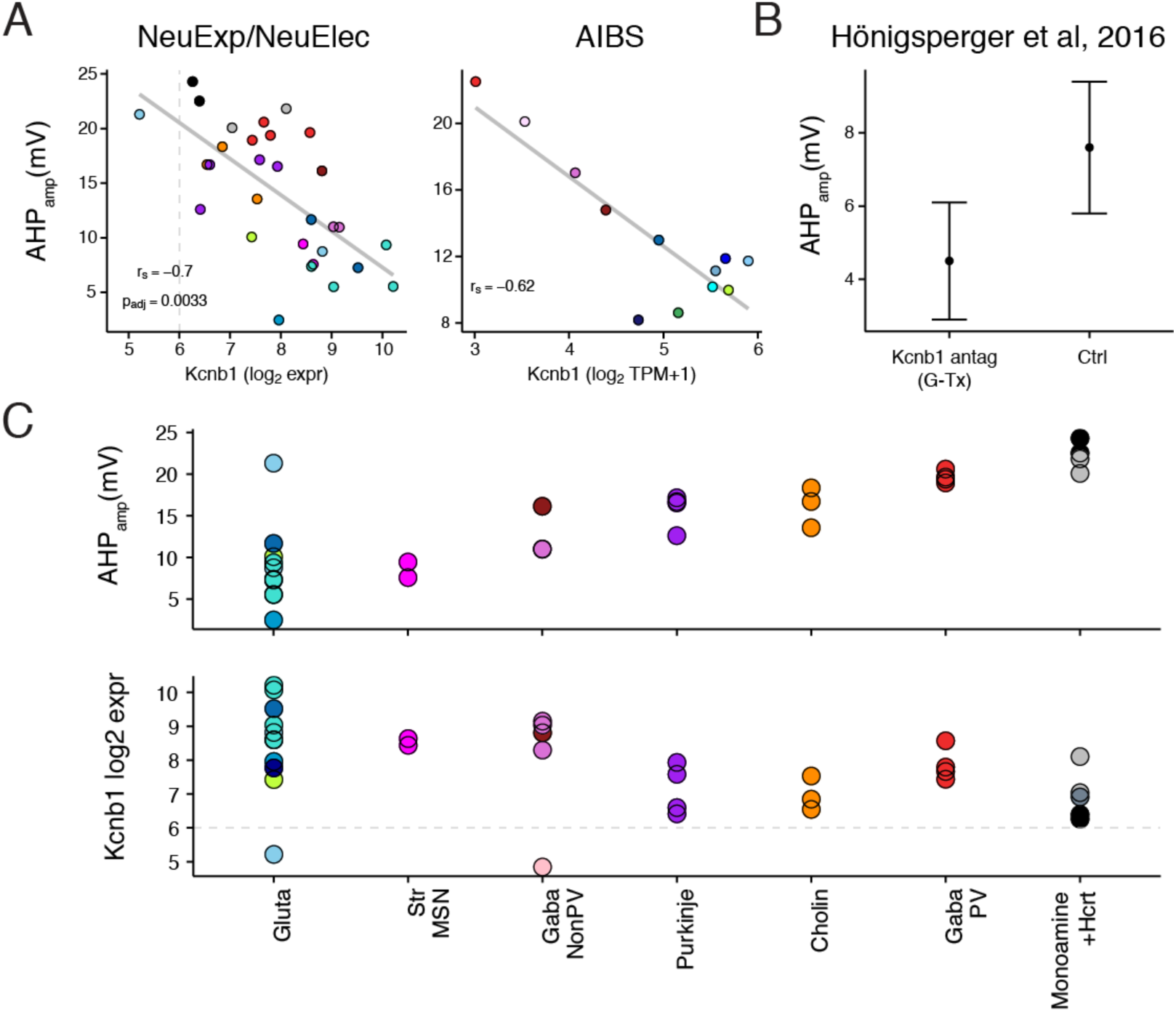
Specific evidence for gene-electrophysiology correlation not implying causation. A) Correlation between cell type-specific *Kcnb1* (K_v_2.1) gene expression and action potential after-hyperpolarization amplitude (AHP_amp_) from discovery dataset (NeuExp/NeuElec, left) and Allen Institute dataset (AIBS, right). B) Replotted data from Honingsperger et al 2016, showing measured AHP_amp_ values from entorhinal cortex pyramidal neurons during control and under perfusion of Guangxitoxin-1E, a specific blocker of K_v_2-family currents. Data illustrates that effect of K_v_2.1 blockade results in increased AHP_amp_, the opposite of expected result based on correlations shown in A. C) Same data shown in A, but broken down by major cell types, illustrating that *Kcnb1*-AHP_amp_ correlation is in part related to major differences in *Kcnb1* expression and AHP_amp_ values between excitatory glutamatergic and non-excitatory cell types.

### Further analysis of specific gene-electrophysiology correlations

Encouraged that many of the ion channel gene-ephys associations discovered through our analysis were consistent with previous experimental manipulations, we next expanded our attention to other classes of genes. From the larger list of correlations identified in our analysis (Supplemental Table 3), we have highlighted below a small number of individual gene-ephys correlations.

Multiple genes known to regulate ion channel functional expression and localization were identified in our analysis (Figure 4A, B). For example, two genes regulating the localization of sodium channels, *L1cam* and *Fgf14*, were correlated with V_rest_ in our analysis and the direction of correlation was further supported by previous experiments ^44,45^. Along this theme, our analysis identified novel associations between *Nedd4l* and *Slmap* with V_rest_, *Ank1* with maximum firing frequency, and *Nkain1* with R_in_ (as shown in Figure 1). *Nedd4l*, identified as an epilepsy gene through whole-exome sequencing ^14^, ubiquitinates voltage-gated sodium and potassium channels ^46^; *Slmap*, associated with Brugada syndrome, controls the trafficking and surface expression of voltage-gated sodium channels in cardiac and muscle cells but remains unstudied in neurons ^47^. *Ank1*, a member of the ankyrin family, has recently been shown to coordinate the localization of specific Na_v_ subunits to nodes of Ranvier ^48^. Though we found the highest expression of *Ank1* in fast-spiking cells, including Purkinje and PV interneurons, its function remains completely uncharacterized in these cells.

We noted several transcription factors in our list of associated genes, including some that have known roles in the nervous system that are compatible with possible, but unknown, roles in the regulation of cellular ephys (*Figure 4*C). For example, we found *Zbtb18* (a.k.a., RP58, Zfp238) to be negatively correlated with V_rest_. Though *Zbtb18* has yet to be studied for its potential electrophysiological effects, this gene has been shown to be required for the normal development of neocortical glutamatergic cells *^49,50^* and its human homolog has recently been identified as a causative gene for autism and neurodevelopmental disorders *^51^*. As another example, *Zscan21* (a.k.a., Zipro1 or Zfp38) positively correlated with input resistance here and has been shown to be involved in the normal proliferation of progenitor cells into cerebellar granule cells *^52^*.

Among genes correlated with membrane capacitance and input resistance, we noticed that many of these were cytoskeletal proteins or otherwise associated with regulating neuronal differentiation and dendritic morphology, including *Cap2*, *Chn1*, and *Bex1*, and *Tpm4* (Supplemental Figure 5).

In summary, this analysis presents suggestive evidence for many novel gene-ephys relationships. Though we do not expect all of these novel associations to reflect direct causal relationships, by focusing on gene classes that are compatible with possible regulation of ephys, we can further hone the list of associated genes to those that might be of further interest for follow-up investigation.

**Figure 4:**
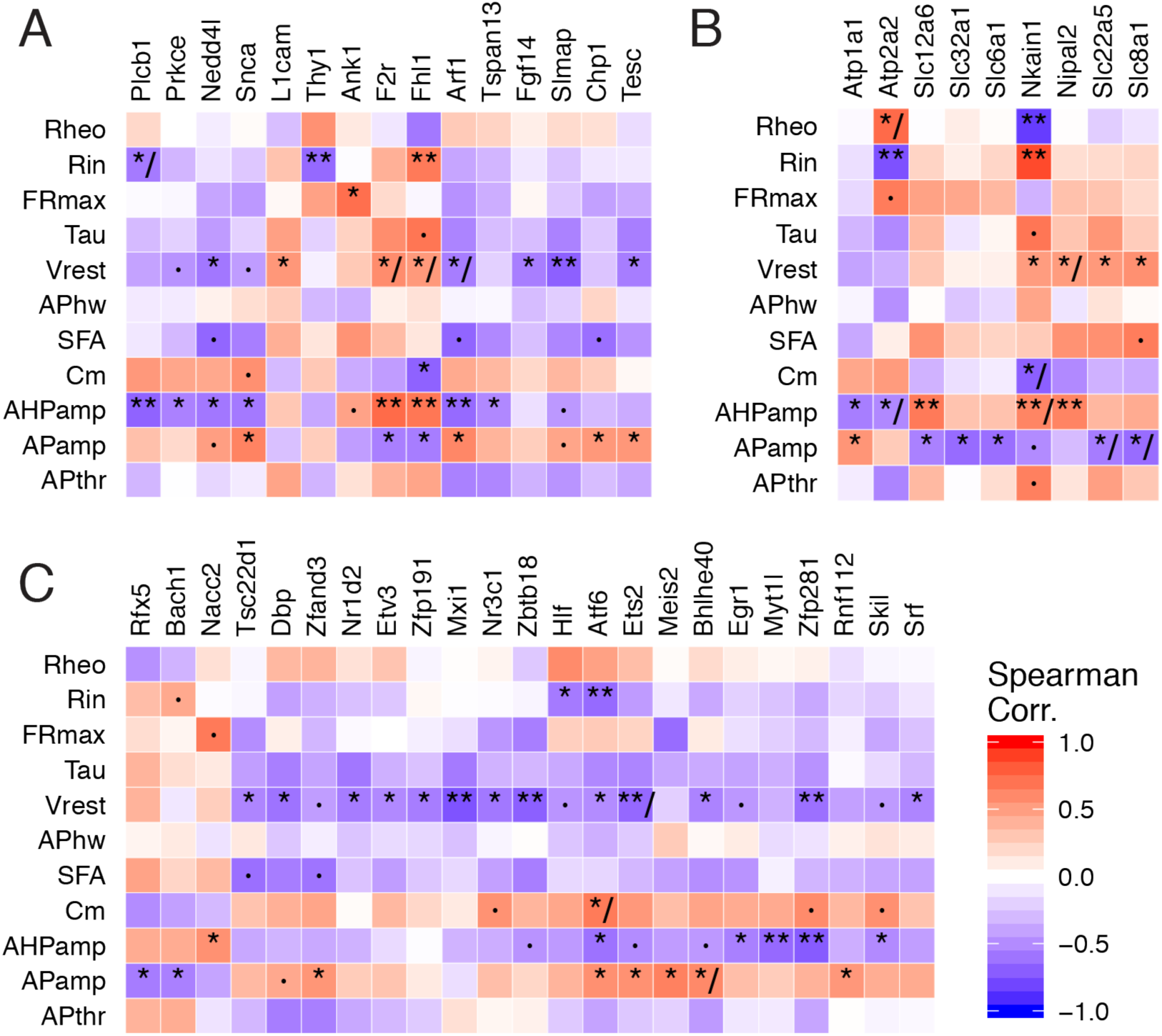
Summary of gene-ephys correlations for selected functional gene sets. A) Genes regulating ion channels and transporter function. B) Ion transporters. C) Transcription factors. Genes filtered for those with at least one statistically significant correlation with an ephys property (p_adj_ < 0.05) and validating in AIBS dataset. Symbols within heatmap: ·, p_adj_ < 0.1; *, p_adj_ < 0.05; **, p_adj_ < 0.01; /, indicates inconsistency between discovery and AIBS dataset.

**Supplemental Figure 5:**
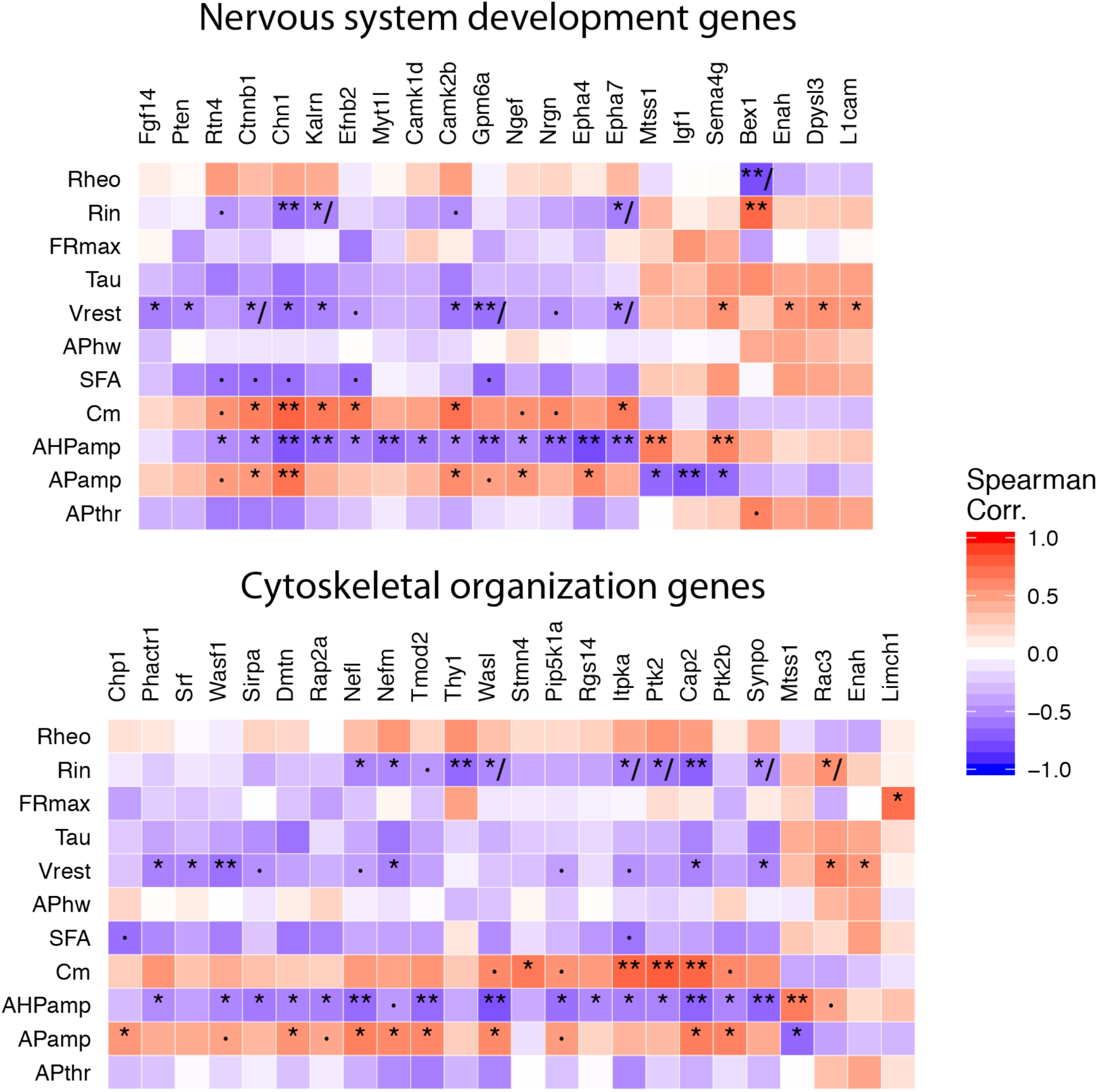
Summary of gene-ephys correlations for additional functional gene sets. *Top: Nervous system development genes. Bottom: Cytoskeletal organization genes. Genes filtered for those with at least one statistically significant correlation with an ephys property (p_adj_ < 0.05) and validating in AIBS dataset. Symbols within heatmap: ·, p_adj_ < 0.1; *, p_adj_ < 0.05; **, p_adj_ < 0.01; /, indicates inconsistency between discovery and AIBS dataset.*

## Discussion

The relationship between gene expression and cellular phenotypes like electrophysiology or morphology is complex and largely unknown. Here, we have enumerated a subset of potential gene-electrophysiology relationships by identifying genes whose expression significantly correlates with specific electrophysiology parameters across a brain-wide collection of neuron types. The majority of these relationships were consistent in an independent sample of visual cortex cell types. Beyond correlation, some of these genes, such as *Scn1a*/Na_v_1.1 and *Gabrd*, have been experimentally shown to be causally responsible for specific ephys properties. The majority of genes discussed here, such as *Nkain1* and *Slmap*, have yet to be investigated in the context of neuronal intrinsic electrophysiology. These genes present opportunities for further study and potential avenues for targeted manipulation of electrophysiological features.

The combined NeuroExpresso/NeuroElectro reference dataset is a first-of-its-kind resource of cell type-specific transcriptomes paired with electrophysiological profiles across a large collection of neuron types. The community resource directly reflects the efforts of hundreds of investigators to characterize the rich diversity of neuron types throughout the brain. It further reflects our considerable efforts in curating and standardizing this heterogeneous data ^23–25^. The dataset includes cell type-specific samples from a wide range of cell types varying in sub-threshold and spiking patterns, morphologies, and developmental stages. We have made the combined dataset available here, as it could be a useful resource and benchmark for future analyses. Moreover, the approach could be expanded to incorporate additional cellular phenotypes, like neuronal morphology or synaptic physiology, and newer genomic data sources including from RNA-seq, epigenomics, or proteomics ^18,53–55^.

In our framework, a causal gene-ephys relationship implies that a consistent change in a gene’s expression would result in a corresponding change in an ephys phenotype, all else being equal. Based on the diversity of cell types present here, we hypothesize that these gene-ephys relationships might further be relatively independent of cell type identity. Indeed, we found examples during our literature search where the specific experiment to confirm a causal gene-ephys relationship was performed in a cell type not present in either the discovery or AIBS datasets, including auditory and autonomic brainstem neurons (Figure 3, Supplemental Figure 4). Not only do these examples provide direct support for the gene-ephys relation, but we also infer the same causal relationship in other cell types, beyond those tested directly. Though additional experiments are needed to determine whether these relationships are truly cell type-independent, this possibility is exciting as it suggests that there could be some genes that contribute to similar ephys functions across very different cell types.

Every novel correlation reported here presents a specific, testable causal prediction. The results from our ion channel-focused literature search are encouraging, as 13 of 17 tested gene-ephys relationships showed some evidence for direct experimental support. However, it is overly optimistic to conclude that most novel ephys-correlated genes reported here will prove causal. Instead, we advocate further in-depth analysis of gene function when prioritizing individual genes for future experiments. For example, the correlation between *Nkain1* and input resistance (R_in_) is plausibly causal because the *Nkain1* protein interacts with the Na^+^/K^+^ pump complex *^29^* and the pump’s activity regulates R_in_ through helping maintain cellular volumes *^30^*. Similarly, the correlation between *Ank1* and FR_max_ is intriguing because *Ank1*, an isoform of the autism gene *Ank3*, helps coordinate the localization of Na_v_ subunits to the nodes of Ranvier *^48^*.

Though we found *Ank1* to be highly expressed in adult PV and Purkinje cells here, its function in these cells has yet to be characterized. Specific transcription factors identified might regulate the expression of downstream genes relevant to ephys. For example, *Zbtb18*, correlated with resting potential here, is required for normal glutamatergic cell development and has recently been implicated in human neurodevelopmental disorders through genome sequencing *^49–51^*. Ultimately, these genes could provide novel means for manipulating cellular ephys in the context of disease. For example, upregulating *Scn1a* expression using anti-sense RNA approaches has been shown to be an effective means of reducing seizures in a model of Dravet syndrome *^32^*.

### Limitations and caveats

The results presented here are restricted to a limited range of situations. First, we can only identify genes where mRNA, as measured in dissociated cells ^56^, is an adequate readout of a gene’s functional activity at the protein level. Future datasets employing RNA-seq, proteomics, or techniques to capture non-somatic mRNA will likely be able to identify more genes where alternative splicing and post-translational modifications are essential for understanding gene function ^10–12^.

Second, our analysis approach assumes a gene’s contribution to electrophysiology is similar and monotonic across cell types. We likely miss genes that contribute to complex ephys features in ways that are biologically degenerate and are highly non-linear or combinatorial ^27,28^. For example, K_v_3-family ion channels, including *Kcnc1*/K_v_3.1, have been implicated in helping fast-spiking cells maintain narrow spike widths ^31,57^, but we did not identify *Kcnc1* as correlated with AP width in our analysis. More sophisticated analysis methods, such as those that incorporate information about how proteins interact to form functional complexes, might reveal additional signals and mitigate spurious correlations. However, pursuing such approaches will likely necessitate larger datasets than are currently available.

Third, the focus of our analysis is to explain how ephys differences across cell types emerge through gene expression. It remains an open question whether the same genes driving large *across* cell type differences would also be the same genes that are defining subtler *within* cell type differences, like amongst olfactory bulb mitral cells or CA1 pyramidal cells ^1,2,53^. As the patch-seq methodology, enabling transcriptomic and ephys characterization from the same single-cell ^19,20^, is further developed and applied, we eagerly anticipate testing these hypotheses. However, small changes in expression of individual genes, as expected within a single cell type, are difficult to reliably detect using current technologies, in part, due to relatively limited sample sizes and technical challenges like “dropouts” ^18^. Indeed, while these patch-seq studies have demonstrated their utility in classifying individual cells into types ^19,20^, how variance in expression of specific genes gives rise to within cell type ephys differences remains largely unaddressed.

Fourth, ephys property correlations and gene co-expression limits the potential specificity of any causal prediction made here. For example, some pairs of ephys properties, like AHP_amp_ and R_in_, are correlated but probably do not share common biophysical underpinnings (Supplemental Figure 2B). Because of this common correlation, genes significantly associated with one ephys feature are more likely to be also associated with other ephys features, potentially spuriously. Similarly, many pairs of genes show correlated expression across samples (i.e., gene co-expression). Gene co-expression often reflects biologically meaningful signals, such as co-regulation by common transcription factors or shared membership in biological pathways and cellular compartments ^58^. However, co-expression makes interpreting individual gene-ephys associations difficult and likely contributes to why we found many more genes for some ephys properties than we would naively expect, such as V_rest_ and AHP_amp_. Future analysis approaches that explicitly consider co-expression might prove useful ^59^.

Lastly, the heterogeneous nature of the compiled NeuroExpresso/NeuroElectro dataset ^23,25,56^ might limit our power to see possible biologically relevant signals and could explain our failure to find genes for some ephys features. For example, because data in NeuroElectro are compiled from different studies collected in the absence of standards for how some ephys properties are defined ^24,60^, this likely limits our downstream attempts at normalization. However, the overall consistency with the AIBS Cell Types dataset, where data were collected using standardized conditions and protocols, suggests that the results shown here are not entirely the result of technical artefacts due to data compilation.

### Future directions

Our findings suggest a number of directions for future study. Can specific gene-ephys relationships be used as biomarkers to detect electrophysiological changes in a disease or treatment context? For example, if *Scn1a*/Na_v_1.1 is upregulated in a cell type, does that serve as a reliable indicator of hyper-excitability? Given the relative ease and growing popularity of single-cell transcriptomics on dissociated cells and nuclei ^18,26^, could these gene-ephys correlations be used to impute ephys phenotypes from transcriptomic signatures alone? Lastly, are the gene-ephys correlations reported here predictive of cell-to-cell variability reported within the same cell type?

In summary, our results suggest that large-scale transcriptomics can prove useful in helping elucidate the biophysical basis for the rich electrophysiological diversity seen amongst neuron types throughout the brain.

## Methods

### NeuroExpresso Database Description

To obtain neuron type-specific transcriptomic data, we made use of the NeuroExpresso database (neuroexpresso.org), described previously ^23^. Briefly, the database contains transcriptomic studies collected from mouse brain cell types sampled under normal conditions. We specifically utilized the microarray-specific subset of NeuroExpresso. These samples were collected using purified, pooled-cell microarrays with transcriptomes quantified using the Affymetrix Mouse Expression 430A Array (GPL339) or Mouse Genome 430 2.0 Array (GPL1261). We further only used probesets that were shared between both platforms. Transcriptomic samples were quality controlled and manually curated for cell type identity and basic sample metadata, including animal age, array platform, and purification method. The samples were subjected to RMA normalization and an additional round of quantile normalization in order to obtain a uniform distribution of signals across samples. When a single gene was represented by multiple probesets, the probeset with highest variability across samples was chosen to represent the gene. We note that we have re-annotated the cell type labels used here from those used in the NeuroExpresso database and web resource.

For the purpose of obtaining a large corpus of cell types, we made use of a small number of cell type-specific transcriptomic samples excluded from analysis in the original NeuroExpresso publication (e.g., developmentally immature samples). Specifically, for two major cell types with transcriptomic data collected at varying ages, cortical parvalbumin-positive (PV) interneurons labelled by the G42 mouse line and cerebellar Purkinje cells ^22,61^, we kept samples collected at different ages separate and used of samples collected from animals aged less than P14. We further included data representing cortical *Htr3a-* and *Oxtr*-expressing cells from Gene Expression Omnibus (GEO) accession GSE56996 ^62^ and layer 2-3 and layer 6 pyramidal cells from GSE69340 ^63^. The complete listing of transcriptomic samples, annotated cell types, and references is provided in Supplemental Table 2.

### Gene filtering and sample summarization

Following data compilation, we filtered genes to retain only those with 1) high mean expression; and 2) highly variable expression across cell types in the combined dataset. Specifically, for each gene, *g*, we calculated its expression mean, μ_g_, and standard deviation, σ_g,_ across the collection of 34 cell types in the combined discovery dataset. Next, we calculated a global mean, μ_global_ defined as *mean*(μ_g1:gN_), and standard deviation, σ_global_ defined as *mean*(σ_g1:gN_) across the total set of genes. Here, μ_global_ = 7.5 and σ_global_ = 0.75; for context, background expression levels were approximately ˜6.0 (log_2_ expression units). We filtered genes where μ_g_ > μ_global_ and σ_g_ > σ_global_, leaving 2694 from 11667 total genes quantified. Lastly, we summarized each cell type by the mean expression per gene across samples.

## NeuroElectro Database Description and Normalization

To obtain neuron type-specific electrophysiological measurements, we used an updated version of the NeuroElectro database (neuroelectro.org), originally described in ^24,25^. Briefly, we populate the NeuroElectro database using manual curation to extract information on electrophysiological measurements such as resting membrane potential and input resistance (described in Supplemental Table 1) from the results sections of published papers using intracellular electrophysiology. Curators also annotate a set of relevant methodological information, including species, animal age, electrode type, preparation type, recording temperature, and use of liquid junction potential correction.

### NeuroElectro database

We note the following major improvements to the NeuroElectro database, beyond an increase in the overall database size (from 331 to 968 articles as of December 2016).

First, we have now curated and manually standardized a greater number of electrophysiological properties, including after hyperpolarization amplitude (AHP_amp_), maximum spiking frequency (FRmax), and spike frequency adaptation (SFA). For example, in the process of data curation we have standardized electrophysiological properties for the use of different baselines, for example, AHP amplitude reported as an absolute voltage as opposed to amplitude relative to spike threshold (e.g., -70 mV vs 10 mV). We note that because of raw data unavailability, we do not recalculate measurements in NeuroElectro from raw ephys traces. Thus, we could not ensure that ephys properties such as SFA or AHP_amp_ were calculated using a consistent stimulation protocol across different studies. These differences where present would tend to contribute to study-to-study variability.

Second, when curating specific neuron subtypes reported in the literature, we now take care to manually annotate the specific features the authors used to define each cell subtype (e.g., the mouse line used, brain region, gene or protein expression, firing pattern, etc.); for example, “barrel cortex layer 2-3 somatostatin-expressing interneuron from the GIN mouse line” or “hypothalamus orexin-expressing cell”. This level of fine-grained cell type curation allows us to better harmonize relevant electrophysiological to transcriptomic datasets post hoc.

### NeuroElectro Data Preprocessing

Electrophysiological data was filtered for: 1) recordings from acute brain slices *in vitro* (thus removing *in vivo* recordings and from slice and cell cultures); *2)* from mice, rats, or guinea pigs; 3) with an animal age greater than 2 days old. Animal ages, when reported as a range (e.g., P14-P20), were summarized using the geometric mean. When animal age or recording temperature was not reported, we used median imputation to fill in missing values (which typically was rare). To address the correction of liquid junction potential (LJP), we manually removed or “uncorrected” the correction of LJP when it had previously been performed and when the original authors provided the explicit voltage correction value used (i.e., LJP offset). We then used a custom LJP metadata field denoted ‘PostCorrected’ to define these cases.

### Experimental condition-based data normalization

As described previously, we used statistical regression models to normalize ephys data for study-to-study differences in experimental methodologies ^25^. Here, we used elastic-net penalized regression, implemented using the *cv.glmnet* function within the R *glmnet* package ^64^ with an alpha value of .99 and nlambda = 100. The regression model for each ephys parameter (EphysProp) was fit using the following formula:

EphysProp = NeuronType + Species + JxnPotential + ElectrodeType + bs(log_10_(AnimalAge)) + bs(RecTemp)

where bs indicates the use of bsplines with 5 degrees of freedom. Here, NeuronType, Species, JxnPotential, and ElectrodeType each indicate nominal metadata types. AnimalAge and RecTemp refer to animal age and slice recording temperature and reflect continuous parameters. For example, ElectrodeType indicates the use of patch-clamp, perforated patch, or sharp electrodes whereas JxnPotential indicates whether the liquid junction potential was explicitly corrected, not corrected, or unmentioned within the article’s methods section. The ephys properties, R_in_, Tau, AP_hw_, C_m_, Rheo, FR_max_, were log_10_-transformed prior to metadata modeling.

We used the filtered NeuroElectro dataset to fit regression models to model study-to-study variability in ephys measurements. After fitting these models, we then used the models to adjust ephys data for the influence of major differences in experimental conditions between studies.

To summarize electrophysiological measurements per each unique cell type (defined in Supplemental Table 2), we first averaged measurements reported within a single paper and then calculated the median ephys value across papers characterizing each cell type.

## Harmonizing cell types across NeuroExpresso and NeuroElectro

Because it was uncommon for a single study to characterize both a cell type’s transcriptomic and electrophysiological parameters, we developed a strategy for pairing gene expression and ephys datasets from different studies based on common cell type identity.

We first manually re-annotated the cell type identity of each transcriptomic sample from NeuroExpresso using a descriptive semantic label (shown in Supplemental Table 2), defined by a minimally sufficient number of defining features (including brain region and marker gene expression or projection pattern ^65^). For example, the transcriptomic samples corresponding to cerebellar granule cells in NeuroExpresso were purified using the L10a-Neurod1 mouse line, where GFP is specifically expressed in the ribosomes of these cells ^66^. Here, we merely annotated these samples using the label, “cerebellar granule cells” (CB gran). We next identified all curated electrophysiological data within NeuroElectro corresponding to this same major cell type, making use of the manual annotations for each electrophysiological sample’s cell type identity (n = 9 articles for CB granule cells). We note that subtle differences between how CB granule cells are labelled in the L10a-Neurod1 mouse line and how CB granule cells are targeted by lamina and morphology for ephys recordings would tend not to be preserved after this data harmonization step.

To pair transcriptomic to ephys datasets explicitly defined by different ages (e.g., P7 and P25), we matched animal ages +/- 2.5 days. For example, the samples corresponding to “Ctx G42 P15” reflect neocortical parvalbumin-positive interneurons labeled by GFP in the G42 mouse line aged P15 +/- 2.5 days. Because we tended to have fewer data points after subsetting the cortical G42 cells into different age groups, for one ephys property, AP_thr_, we excluded AP_thr_ values from these cells since they varied widely (˜10mV) across studies from the same time point.

## Allen Institute for Brain Sciences Cell Types dataset

### Single cell transcriptomic samples

We made use of an Allen Institute for Brain Sciences (AIBS) Cell Types dataset employing single-cell RNAseq to characterize diversity of cells in adult mouse visual cortex labelled by different mouse cre-lines. Specifically, we obtained data originally reported in ^26^ from GSE71585, representing data from 1809 single-cells. We made use of the summary data file where expression for each gene was summarized as reads per kilobase sequenced per million (TPM) with 24,057 genes quantified per cell.

### Single cell electrophysiological samples

We made use of the AIBS Cell Types dataset employing *in vitro* patch clamp electrophysiology to characterize mouse visual cortex cellular intrinsic electrophysiology using standardized protocols. For each cell in the AIBS Cell Types database (http://celltypes.brain-map.org/), representing 847 single cells as of December 2016, we downloaded its corresponding raw and summarized ephys data (summary measurements included input resistance and resting potential). For all spiking measurements except maximum firing rate and spike frequency adaptation, we used the voltage trace corresponding to the first spike at rheobase stimulation level. For a few ephys properties, like action potential half width, we calculated these from the raw ephys traces, as these were not available in the pre-calculated summarized data. Membrane capacitance was defined as the ratio of the membrane time constant to the membrane input resistance. Maximum firing rate and spike frequency adaptation were calculated using the voltage trace corresponding to the current injection eliciting the greatest number of spikes. Spike frequency adaptation (SFA) was defined as the ratio between the first and mean inter-spike intervals during this maximum spike-eliciting trace (i.e., neurons with greater SFA will show values closer to 0).

### Data summarization and harmonization

We summarized single cell transcriptomic and ephys data to the level of cell types by averaging measurements within the same cre-line (i.e., defining cell types by unique cre-lines). We filtered cre-lines that were sampled by at least 10 cells in each of the transcriptomic and ephys data, leaving a total of 12 cell types / cre-lines. We also filtered single cell transcriptomic samples to include only those corresponding to neuronal cells (i.e., removing glial cells erroneously labelled by the cre-line). We did not further attempt to make use of the novel transcriptomics-based cellular subtypes as defined in ^26^, since we cannot make a correspondence between these subtypes (defined on the basis of multivariate gene expression in the absence of ephys or morphological characterization) with individual cells sampled in the ephys data. We matched genes across the AIBS and NeuroExpresso/NeuroElectro datasets using NCBI entrez gene identifiers. Of the total 2694 genes present in the discovery dataset after expression level-based filtering, there were 2603 total genes in common with the AIBS scRNAseq dataset.

## Data availability

The harmonized and processed cell type-specific data for the discovery and validation datasets has been made publically available at http://hdl.handle.net/11272/10485.

## Statistical analysis and methodology

### Gene-electrophysiological property correlation analysis

For each gene in the filtered NeuroExpresso/NeuroElectro data matrix, we calculated its rank correlation and uncorrected p-value (two-sided test) with each the 11 ephys properties, using the function *cor.test* from the R *stats* package, with ‘method=”spearman”’. We also calculated the Spearman correlation (r_s_) for each gene and ephys property in the AIBS validation dataset.

### Corrections for multiple comparisons

We used the Benjamini-Hochberg correction for False Discovery Rate (FDR) to correct for comparisons performed across multiple genes^67^, implemented using the function *p.adjust* from the R *stats* package. Here, for ease of interpretation, we refer to the Benjamini-Hochberg FDR as p_adj_. Because of ephys property correlations, we did not further correct for multiple comparisons across ephys properties.

### Comparing results across discovery and validation datasets

To evaluate the consistency between discovery and validation datasets, we defined two separate measures. First, to obtain a measure of the overall consistency per ephys property, we calculated the rank correlation across the set of 2603 genes in common to both datasets (after filtering genes for expression levels based on the discovery dataset). Second, to specifically focus on gene-ephys correlations meeting our threshold for significance in the discovery dataset (p_adj_ < 0.05), we defined consistent correlations as those with matching correlation directions and also with the absolute value of the gene-ephys rank correlation in the validation dataset exceeding 0.3 (i.e., |r_s, validation_| > 0.3). For both criteria, we obtained p-values through randomly shuffling cell type labels in the validation dataset between ephys and gene expression data. We obtained an expected p-value null distribution through performing 1000 random shuffles and recalculating gene-ephys correlations per shuffle. Our final list of gene-ephys correlations are those that are significant in the discovery dataset (i.e., p_adj, discovery_ < 0.05) that further validated in the AIBS dataset (|r_s, validation_| > 0.3).

## Gene lists

To obtain specific gene sets, we made use of Gene Ontology annotations (as of August 2016). We used the GO term 0005216 corresponding to “ion channel activity” to identify ion channels; the term 0015075 corresponding to “ion transmembrane transporter activity” in addition to *Nkain1* to identify ion transporters; the term 0007010 corresponding to “cytoskeleton organization” to identify cytoskeletal genes; the term 0007399 corresponding to “nervous system development” to identify developmental genes; and the term 0034765 to identify “regulation of ion transport” in addition to the genes *L1cam, Slmap,* and *Ank1*. To obtain a comprehensive manually curated listing of transcription factors, we used the Transcription Factor Checkpoint resource ^68^.

## Ion channel focused literature search

### Literature search methodology

We performed a systematic literature search to identify causal experiments consistent or inconsistent with the individual gene-ephys correlations reported here. Specifically, we started with a set of 23 ion channel genes identified by our analysis (defined by GO term 0005216) that further validated in the AIBS dataset.

For each gene, we manually searched for articles where these genes had been perturbed, either using genetic approaches to knockout or knockdown the gene’s expression or using channel-specific pharmacology. When searching for individual genes, we made use of common gene name synonyms, for example, that K_v_1.1 is a synonym for the gene *Kcna1*. We further searched for papers where the individual ephys properties suggested by our correlative analysis (e.g., AP_hw_, rheobase) had been explicitly measured. To this end, we used Google Scholar with the gene name or gene name synonym and the associated ephys property as search terms. When the name of a pharmacological blocker of an ion channel was known it was included in search terms. We also checked the top 40 papers related to a gene on its NCBI Gene page for those in which the gene was manipulated and ephys properties of interest were measured. For some widely studied ion channel genes, such as *Kcna1*/K_v_1.1 and *Kcnd2*/K_v_4.2, we did not attempt to systematically review each article studying these genes and typically ended our search after 3-5 relevant articles were identified. We further limited our assessment to perturbations involving mammalian neurons.

When our search yielded pertinent articles, we annotated relevant information, including: the kind of manipulation (e.g., genetic manipulation and type; pharmacological compound used, etc.); cell type; and direction and magnitude of effect. To categorize effects, we assessed whether the perturbation resulted in an increase or decrease in the value of the ephys property and whether this change was further either statistically significant or non-significant. In a small number of cases, there was effectively no change or a negligible change between the control and perturbed condition that were curated as “negligible changes”.

When scoring whether an individual gene-ephys correlation was either consistent or inconsistent with literature evidence, we assessed the direction effect. For example, for an ion channel gene that our analysis found as positively correlated with V_rest_, we would expect that knocking out the gene would make V_rest_ to become more negative and more hyperpolarized, all else being equal. Similarly, applying an agonist of the ion channel should make V_rest_ more positive and depolarized. In cases with multiple lines of evidence linking specific ion channel perturbations to ephys changes (e.g., both pharmacological and genetic changes), we aggregated these along the following categories: consistent, inconsistent, mixed, and no effect. Gene-ephys correlations supported by both consistent and inconsistent literature evidence were marked as “mixed”. Those with consistent evidence and also some evidence for a negligible change but no inconsistent evidence were marked as “consistent”, and similarly for inconsistent evidence.

## Acknowledgments

We thank the Pavlidis Lab undergraduates for assistance with database curation. We thank R. Richardet and S. Hill for aid with cell type ontologies. We thank Steve Prescott, Jesse Gillis, Megan Crow, and members of the Pavlidis Lab for helpful discussions and comments on the manuscript. We are especially grateful to all of the investigators whose data are represented in the NeuroExpresso, NeuroElectro, and Allen Institute for Brain Sciences Cell Types databases. This work is supported by a NeuroDevNet grant to PP, the UBC bioinformatics graduate training program to BOM, a CIHR post-doctoral fellowship to SJT, a NSERC Discovery grant (RGPIN-2016-05991) and NIH grants MH111099 and GM076990 to PP.

## Supplemental Tables

**Supplemental Table 1.**
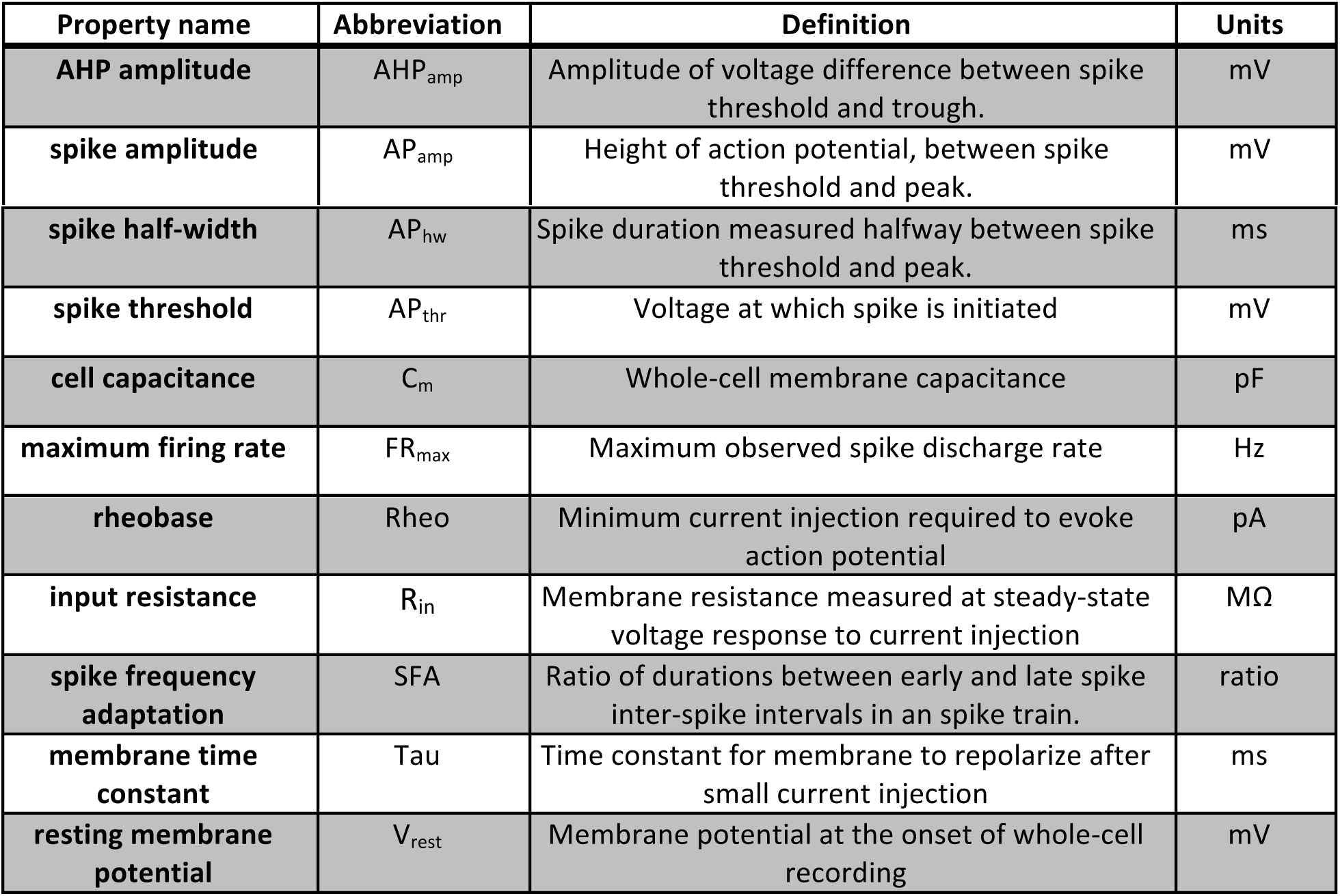
Description of electrophysiological properties used in this study.

| Property name | Abbreviation | Definition | Units |
| --- | --- | --- | --- |
| **AHP amplitude** | AHP<sub>amp</sub> | Amplitude of voltage difference between spike threshold and trough. | mV |
| **spike amplitude** | AP<sub>amp</sub> | Height of action potential, between spike threshold and peak. | mV |
| **spike half-width** | $AP_{hw}$ | Spike duration measured halfway between spike threshold and peak. | ms |
| **spike threshold** | $AP_{thr}$ | Voltage at which spike is initiated | mV |
| **cell capacitance** | $C_m$ | Whole-cell membrane capacitance | pF |
| **maximum firing rate** | $FR_{max}$ | Maximum observed spike discharge rate | Hz |
| **rheobase** | Rheo | Minimum current injection required to evoke action potential | pA |
| **input resistance** | $R_{in}$ | Membrane resistance measured at steady-state voltage response to current injection | $M\Omega$ |
| **spike frequency adaptation** | SFA | Ratio of durations between early and late spike inter-spike intervals in an spike train. | ratio |
| **membrane time constant** | Tau | Time constant for membrane to repolarize after small current injection | ms |
| **resting membrane potential** | $V_{rest}$ | Membrane potential at the onset of whole-cell recording | mV |

**Supplemental Table 2:** Description of cell types composing the combined NeuroExpresso/NeuroElectro dataset.

| **Abbreviation** | **Neuron Type** | **Neuron Matching Criteria** | **Expression References** | **Ephys References** |
| --- | --- | --- | --- | --- |
| BF ACh | Basal forebrain cholinergic cells | in:basal forebrain, neurotransmitter:cholinergic | Doyle et al., 2008 | Matthews et al. (1999), Bengtson et al. (2000), Henderson et al. (2004), Arrigoni et al. (2006), Hedrick et al. (2010), Kalmbach et al. (2012), Unal et al. (2012), McKenna et al. (2013), Yi et al. (2015), Dannenberg et al. (2015), Rosati et al. (1999) |
| BLA Pyr | Basolateral amygdala pyramidal cells | in:basolateral amygdala, morphology:pyramidal | Sugino et al., 2006 | Wu et al. (2007), Kaneko et al. (2008), Jasnow et al. (2009), Ehrlich et al. (2012), Daftary et al. (2012), Motanis et al. (2014), Ferreira et al. (2015), Maiya et al. (2015), Rau et al. (2015) |
| BS ACh | Brain stem cholinergic cells | in:brainstem, neurotransmitter:cholinergic | Doyle et al., 2008 | Brown et al. (2006), Leijon et al. (2014), Kamii et al. (2015) |
| CB Golgi | Cerebellum Golgi cells | in:Cerebellum, morphology:Golgi | Doyle et al., 2008 | Robberechts et al. (2010), Hirono et al. (2012) |
| CB gran | Cerebellum granule | in:Cerebellum, | Doyle et al., 2008 | Brickley et al. (2001), Cathala et |
|  | cells | morphology:granule |  | al. (2003), Gall et al. (2003), Goldfarb et al. (2007), Prestori et al. (2008), Osorio et al. (2010), Usowicz et al. (2012), Delvendahl et al. (2015), D'Angelo et al. (1998) |
| CB Purk P14 | Cerebellum Purkinje cells, P14 | in:Cerebellum, morphology:Purkinje, age: P14 | Paul et al 2012 | Swensen et al. (2005), McKay et al. (2005), Zhu et al. (2006), Takeuchi et al. (2008) |
| CB Purk P3 | Cerebellum Purkinje cells, P3 | in:Cerebellum, morphology:Purkinje, age: P3 | Paul et al 2012 | McKay et al. (2005) |
| CB Purk P56 | Cerebellum Purkinje cells, P56 | in:Cerebellum, morphology:Purkinje, age: P56 | Paul et al 2012 | McKay et al. (2005), Torashima et al. (2006), Hoxha et al. (2012) |
| CB Purk P7 | Cerebellum Purkinje cells, P7 | in:Cerebellum, morphology:Purkinje, age: P7 | Paul et al 2012 | McKay et al. (2005) |
| DG gran | Dentate gyrus granule cells | in:Dentate gyrus, morphology:granule | Perrone-Bizzozero NI et al. 2011 | Scharfman et al. (2000), Joëls et al. (2001), Romo-Parra et al. (2003), Kobayashi et al. (2003), Scharfman et al. (2003), Patel et al. (2004), Dietrich et al. (2005), Podlogar et al. (2006), Patrylo et al. (2007), Zhan et al. (2009), Epszstein et al. (2010), Powell et al. (2012), Zhang et al. (2012), Kim et al. (2012), Swijsen et al. (2012), Lopez et al. (2012), Cabezas et al. (2013), Dieni et al. (2013), Yarishkin et al. (2014), Nenov et al. (2015), Pourbadie et al. (2015), Gao et al. (2015), Mott et al. (1997), Lübke et al. (1998) |
| ORB L5 Pyr | Frontal cortex layer 5 pyramidal cells | in:frontal cortex, morphology:pyramidal, layer:layer 5 | Sugino et al., 2006 | Lavin et al. (2001), Noaín et al. (2006), Orozco-Cabal et al. (2008), Zhong et al. (2008), Gullledge et al. (2009), van et al. (2009), Yan et al. (2011), Winters et al. (2011), Hirai et al. (2012), Sepulveda-Orengo et al. (2013), Wang et al. (2013), van et al. (2015), Yang et al. (2013), Zhong et al. (2014), Stephens et al. |
|  |  |  |  | (2014), Sceniak et al. (2016), Song et al. (2015), de et al. (1996) |
| CA1 Pyr | Hippocampus CA1 pyramidal cells | in:hippocampus CA1, morphology:pyramidal | Sugino et al., 2006 | Karnup et al. (1999), Kirson et al. (2000), Beier et al. (2000), Jahromi et al. (2000), Staff et al. (2000), Pike et al. (2000), Fraser et al. (2001), Sanabria et al. (2001), Foster et al. (2001), Su et al. (2001), Gusev et al. (2001), Thibault et al. (2001), Ireland et al. (2002), Castro et al. (2002), Vreugdenhil et al. (2002), Martina et al. (2003), Kamal et al. (2003), Oh et al. (2003), Scammell et al. (2003), Carrer et al. (2003), van et al. (2003), McDermott et al. (2003), Li et al. (2004), Fujiwara-Tsukamoto et al. (2004), Yue et al. (2004), Barbaro et al. (2004), Martina et al. (2005), Golding et al. (2005), McCloskey et al. (2005), Yue et al. (2005), Avignone et al. (2005), Otto et al. (2006), Grabauskas et al. (2006), Pandis et al. (2006), Gant et al. (2006), Cao et al. (2006), Shelbourne et al. (2007), Lamsa et al. (2007), Derchansky et al. (2008), López et al. (2007), Yang et al. (2008), Yan et al. (2008), Pilpel et al. (2009), Maggio et al. (2009), Lopantsev et al. (2009), Routh et al. (2009), Patino et al. (2009), Tartar et al. (2010), Gamelli et al. (2011), Zemankovics et al. (2010), Chevaleyre et al. (2010), Roggenhofer et al. (2010), Dasari et al. (2011), Scorza et al. (2011), Gant et al. (2011), Sinning et al. (2011), Drew et al. (2011), Edelmann et al. (2011), Kaphzan et al. (2011), Malik et al. (2012), Zhou et al. (2011), None et al. (2012), Farmer et al. (2012), Marcelin et al. (2012), Kim et al. |
|  |  |  |  | (2012), Graves et al. (2012), McLeod et al. (2013), McKay et al. (2013), Oh et al. (2013), San et al. (2014), Borel et al. (2013), Booth et al. (2014), Groen et al. (2014), Pillai et al. (2014), Johnson et al. (2014), Springer et al. (2014), Novkovic et al. (2015), Hall et al. (2015), Hooper et al. (2016), Royeck et al. (2015), Cembrowski et al. (2016), Hackert et al. (2016), Fraser et al. (1996), Spigelman et al. (1996), Baraban et al. (1997), Miller et al. (1997), Rempe et al. (1997), Vida et al. (1998), Gulyás et al. (1998) |
| HIP GIN | Hippocampus GIN (SST) interneurons | in:hippocampus, mouse_line:GIN | Sugino et al., 2006 | Griguoli et al. (2009), Maisano et al. (2012), Kim et al. (2012), McKay et al. (2013) |
| HY orexin | Hypothalamus hypocretinergic cells | in:Hypothalamus, expresses:Hcrt | Dalal et al 2013 | Yamanaka et al. (2010), Schöne et al. (2011), Tsunematsu et al. (2011) |
| LC NAdr | Locus cereuleus noradrenergic cells | in:Locus Cereuleus, neurotransmitter:noradrenaline | Sugino et al. 2014 | Jedema et al. (2004), Chang et al. (2006), Taneja et al. (2009), Zhang et al. (2010), de et al. (2010), de et al. (2011) |
| MB 5HT | Midbrain serotonergic cells | in:midbrain, neurotransmitter:serotonin | Dougherty et al 2013 | Li et al. (2001), Kirby et al. (2003), Beck et al. (2004), Crawford et al. (2010), Shikanai et al. (2012) |
| Ctx CStr Pyr | Neocortex corticostriatal pyramidal cells | in:neocortex, morphology:pyramidal, projection:corticostriatal | Schmidt et al 2012 | Hattox et al. (2007), Groh et al. (2010), Suter et al. (2013), Oswald et al. (2013), Guan et al. (2015) |
| Ctx CThal Pyr | Neocortex corticothalamic pyramidal cells | in:neocortex, morphology:pyramidal, projection:corticothalamic | Sugino et al., 2006 | Hattox et al. (2007), Kumar et al. (2008), Llano et al. (2009), Tanaka et al. (2011), Hirai et al. (2012), Oswald et al. (2013) |
| Ctx G42 P10 | Neocortex G42 (PV) interneurons, P10 | in:neocortex, mouse_line:G42, age: | Okaty et al., 2009 | Okaty et al. (2009), Oswald et al. (2011), Pangratz-Fuehrer et al. |
|  |  | P10 |  | (2011), Yang et al. (2014) |
| Ctx G42<br>P15 | Neocortex G42 (PV)<br>interneurons, P15 | in:neocortex,<br>mouse_line:G42, age:<br>P15 | Okaty et al., 2009 | Okaty et al. (2009), Oswald et al.<br>(2011), Pangratz-Fuehrer et al.<br>(2011), Yang et al. (2014), Sippy<br>et al. (2013) |
| Ctx G42<br>P25 | Neocortex G42 (PV)<br>interneurons, P25 | in:neocortex,<br>mouse_line:G42, age:<br>P25 | Okaty et al., 2009 | Okaty et al. (2009), Oswald et al.<br>(2011), Yang et al. (2014) |
| Ctx G42<br>P7 | Neocortex G42 (PV)<br>interneurons, P7 | in:neocortex,<br>mouse_line:G42, age:<br>P7 | Okaty et al., 2009 | Okaty et al. (2009), Pangratz-Fuehrer et al. (2011), Yang et al.<br>(2014) |
| Ctx GIN | Neocortex GIN<br>(SST) interneurons | in:neocortex,<br>mouse_line:GIN | Sugino et al.,<br>2006 | Ma et al. (2006), Halabisky et al.<br>(2006), Xu et al. (2006),<br>Fanselow et al. (2010), Fino et al.<br>(2011), Levy et al. (2012),<br>Kinnischtzke et al. (2012), Sippy<br>et al. (2013) |
| Ctx Glt<br>Pyr | Neocortex Glt25d2-expressing<br>pyramidal cells | in:neocortex,<br>morphology:pyramid<br>al, expresses:Glt25d2 | Schmidt et al<br>2012 | Groh et al. (2010), Guan et al.<br>(2015), Kim et al. (2015) |
| Ctx<br>Htr3a | Neocortex Htr3a-expressing cells | in:neocortex,<br>expresses:Htr3a | Nakanima et. Al<br>2014 | Lee et al. (2010) |
| Ctx L2-3 Pyr | Neocortex layer 2-3<br>pyramidal cells | in:neocortex,<br>morphology:pyramid<br>al, layer:layer 2-3 | Shrestha et al.<br>2015 | Sutor et al. (2000), Aramakis et<br>al. (2000), Lambe et al. (2000),<br>van et al. (2000), Castro et al.<br>(2002), Karpuk et al. (2003),<br>Telfeian et al. (2003), Abel et al.<br>(2004), Bender et al. (2003),<br>Tateno et al. (2004),<br>Hadjilambreva et al. (2005), Lee<br>et al. (2005), Povysheva et al.<br>(2006), Guan et al. (2007),<br>Cheetham et al. (2007), Lee et al.<br>(2007), Lemtiri-Chlieh et al.<br>(2007), Kobayashi et al. (2008),<br>Shruti et al. (2008), Santini et al.<br>(2008), Oswald et al. (2008),<br>Andjelic et al. (2009), Lefort et<br>al. (2009), Gullledge et al. (2009),<br>Cummings et al. (2009), Brill et<br>al. (2010), Cho et al. (2010),<br>Takei et al. (2010), Tanaka et al.<br>(2011), Hirai et al. (2012), Levy<br>et al. (2012), Kinnischtzke et al. |
|  |  |  |  | (2012), Cruikshank et al. (2012), Otis et al. (2013), Sippy et al. (2013), van et al. (2015), Yang et al. (2013), Pillai et al. (2014), Cruz et al. (2014), Rebello et al. (2014), Guan et al. (2015), Neske et al. (2015), Tyler et al. (2015), Ferreira et al. (2015), Barnes et al. (2015), Lacroix et al. (2015), Yang et al. (1997), Pineda et al. (1998) |
| Ctx L6 Pyr | Neocortex layer 6 pyramidal cells | in:neocortex, morphology:pyramidal, layer:layer 6 | Shrestha et al. 2015 | Lambe et al. (2000), Lavin et al. (2001), Brumberg et al. (2003), Karayannis et al. (2007), Lee et al. (2007), Kumar et al. (2008), Lefort et al. (2009), Llano et al. (2009), Ledergerber et al. (2010), Winters et al. (2011), Tanaka et al. (2011), Marx et al. (2013), van et al. (2015), Yang et al. (2013), Tian et al. (2014), Proulx et al. (2015), de et al. (1996), Yang et al. (1997), Xiang et al. (1998) |
| Ctx Oxtr | Neocortex Oxtr-expressing cells | in:neocortex, expresses:Oxtr | Nakanima et. Al 2014 | Nakajima et al. (2014) |
| SSp TT Pyr | Somatosensory cortex layer 5 pyramidal cells | in:somatosensory cortex, morphology:thick tufted, morphology:pyramidal | Sugino et al., 2006 | Le et al. (2007), Hattox et al. (2007), Groh et al. (2010), Suter et al. (2013), Guan et al. (2015), Harb et al. (2016) |
| Str ACh | Striatum cholinergic cells | in:striatum, neurotransmitter:cholinergic | Doyle et al., 2008 | Bennett et al. (2000), Lin et al. (2003), Fino et al. (2008), Gittis et al. (2010), Sanchez et al. (2011) |
| Str Drd1 MSN | Striatum Drd1-expressing medium spiny neurons | in:striatum, expresses:Drd1 | Tan et al 2013, Maze et al 2014, Heiman et al 2014, Doyle et al., 2008 | Cepeda et al. (2008), Gertler et al. (2008), Planert et al. (2013) |
| Str Drd2 MSN | Striatum Drd2-expressing medium spiny neurons | in:striatum, expresses:Drd2 | Maze et al 2014, Heiman et al 2014, Doyle et al., 2008 | Cepeda et al. (2008), Gertler et al. (2008), Planert et al. (2013) |
| SNc DA | Substantia nigra | in:substantia nigra | Chung et al., | Nedergaard et al. (1999), |
|  | pars compacta dopaminergic cells | pars compacta, neurotransmitter:dopamine | 2005 | Neuhoff et al. (2002), Saitoh et al. (2004), Foehring et al. (2009), Seutin et al. (2010), Tateno et al. (2011), Gantz et al. (2011), Ding et al. (2011), Tucker et al. (2012), Branch et al. (2014), Dufour et al. (2014), Richards et al. (1997) |
| VTA DA | Ventral tegmental area dopaminergic cells | in:ventral tegmental area, neurotransmitter:dopamine | Chung et al., 2005 | Neuhoff et al. (2002), Saitoh et al. (2004), Mathon et al. (2005), Smits et al. (2005), Koyama et al. (2006), Margolis et al. (2006), Tateno et al. (2011), Hnasko et al. (2012), Liu et al. (2014) |

*Supplemental Table 3 Legend (Table provided with Supplementary Materials): List of significant gene-electrophysiological correlations.* Column headers are as follows: DiscCorr refers to the gene-ephys rank correlation calculated in the NeuroExpresso/NeuroElectro discovery dataset and DiscFDR and DiscUncorrPval refers to the Benjamini-Hochberg FDR and uncorrected p-value based on this correlation. AIBSCorr refers to the gene-ephys rank correlation in the AIBS replication sample and AIBSConsistent refers to consistency of correlation direction between the discovery and replication datasets.

*Supplemental Table 4 Legend (Table provided with Supplementary Materials): Complete dataset of literature search for ion channels predicted to be significantly correlated with electrophysiological diversity.*

